# The hit-and-run of cell wall synthesis: LpoB transiently binds and activates PBP1b through a conserved allosteric switch

**DOI:** 10.1101/2024.12.13.628440

**Authors:** Irina Shlosman, Andrea Vettiger, Thomas G. Bernhardt, Andrew C. Kruse, Joseph J. Loparo

## Abstract

The peptidoglycan (PG) cell wall is the primary protective layer of bacteria, making the process of PG synthesis a key antibiotic target. Class A penicillin-binding proteins (aPBPs) are a family of conserved and ubiquitous PG synthases that fortify and repair the PG matrix. In gram-negative bacteria, these enzymes are regulated by outer-membrane tethered lipoproteins. However, the molecular mechanism by which lipoproteins coordinate the spatial recruitment and enzymatic activation of aPBPs remains unclear. Here we use single-molecule FRET and single-particle tracking in E. coli to show that a prototypical lipoprotein activator LpoB triggers site-specific PG synthesis by PBP1b through conformational rearrangements. Once synthesis is initiated, LpoB affinity for PBP1b dramatically decreases and it dissociates from the synthesizing enzyme. Our results suggest that transient allosteric coupling between PBP1b and LpoB directs PG synthesis to areas of low peptidoglycan density, while simultaneously facilitating efficient lipoprotein redistribution to other sites in need of fortification.

## Introduction

Penicillin-binding proteins (PBPs) are peptidoglycan (PG) synthases that are among the earliest identified and most successful targets for antibiotic development^1–3^. Penicillin and the related beta-lactam drugs inhibit the enzymatic activity of PBPs^4,5^. However, instances of antibiotic resistance have now been reported for every beta-lactam antibiotic in clinical use, highlighting the need for development of new strategies to target PBP activity^4–10^. A better understanding of the regulatory pathways that control cell wall synthesis by these enzymes will enable such discovery efforts.

PBPs synthesize peptidoglycan (PG) from the precursor lipid II in two enzymatic steps. The disaccharide monomer unit of lipid II is first polymerized by glycosyltransferases (GT) into glycan strands; these nascent strands are then crosslinked by transpeptidases to the existing PG mesh to expand the wall^11–14^. In virtually all bacteria, polymerization and crosslinking reactions are facilitated by two distinct classes of PBP synthases. Class A PBPs (aPBPs) are bifunctional enzymes that possess both GT and TP activities^15^, whereas class B PBPs (bPBPs) are monofunctional transpeptidases that require a glycosyltransferase partner from the SEDS (shape, elongation, division, sporulation) family to carry out synthesis^16–20^. *E. coli* encodes two major aPBPs, PBP1a and PBP1b, and two essential SEDS-bPBP complexes, RodA-PBP2 and FtsW-FtsI (PBP3)^15,16,21–23^. Despite being enzymatically equivalent, these synthetic machines are functionally and structurally distinct and mediate different physiological processes in cells. RodA-PBP2 and FtsW-FtsI synthases associate with multi-protein assemblies to accomplish directed PG synthesis during bacterial elongation and division^21,24,25^.

PBP1a and PBP1b, on the other hand, are thought to fortify and repair the ordered PG framework produced by the SEDS-bPBP complexes^21,26,27^. Accordingly, deletion or inactivation of both PBP1a and PBP1b leads to cell wall damage-induced bulging and lysis^27–30^.

*E. coli* PBP1a and PBP1b are regulated by the outer-membrane tethered lipoproteins (Lpo) LpoA and LpoB, respectively^30,31^. *In vitro* these accessory factors bind dedicated domains on their cognate synthases and accelerate their GTase and TPase activities^32–37^. In a cellular context, Lpo proteins must penetrate through gaps in PG to reach aPBPs in the periplasm, suggesting a mechanism for cell wall damage sensing, whereby lipoproteins preferentially recruit aPBPs to sites of low peptidoglycan density to reinforce the PG matrix^27,34^. However, it has remained unclear how lipoprotein binding is coupled to enzymatic activation, as well as how Lpo-aPBP complexes are disassembled at the end of synthesis, given that the process of repair is expected to seal off PG passageways that lipoproteins rely on for their redistribution.

We previously demonstrated that enzymatic activation of both GT and TP reactions in *Ec*RodA-PBP2 is controlled through structural rearrangements at the GT-TP interface of the enzymatic complex^38^. Mutations in PBP2 that accelerate structural dynamics lead to cell lengthening and allow the synthase to bypass the requirement for cellular activators^38,39^. Notably, suppressor mutations that bypass the deletion of *lpoA* or *lpoB* also map to the interface between the GT and TP domains of their cognate aPBPs^32,33^, suggesting that aPBPs may rely on similar allosteric changes to control their cellular activity. Structural studies of *Ec*PBP1b show that it can adopt distinct conformational states, lending further support to this hypothesis^40–42^.

Here we leveraged single-molecule FRET (smFRET) to investigate if aPBPs undergo conformational rearrangements as part of their catalytic cycle and how these putative rearrangements are controlled by lipoprotein activators. Using the *E. coli* PBP1b-LpoB complex as a model system, we show that LpoB binding stimulates structural changes at the GT-TP interface of PBP1b that trigger enzymatic activation. Following glycan synthesis initiation, the stability of the PBP1b-LpoB complex decreases dramatically, suggesting a mechanism for LpoB recycling. Collectively, our results support a model in which allosteric regulation of PBP1b by LpoB ensures precise targeting of PBP1b synthesis activity to sites in need of fortification or repair, while also optimizing the number of sites that can be fortified through rapid redistribution of LpoB.

## Results

### *E. coli* PBP1b adopts distinct active and inactive states

To test whether structural dynamics contribute to PBP1b function, we developed a single-molecule FRET assay to monitor rearrangements between the GT and TP enzymatic domains of the protein. To this end, we introduced two cysteines (PBP1b^E187C*-*R300C^) into the background of a functional truncation of PBP1b^40,41^ encompassing residues 58-804 (**Fig. 1a**, also *Methods*) and confirmed that the resulting construct (WT) was monodispersed on size-exclusion chromatography, labeled specifically with Cy3- and Cy5-fluorophores and retained enzymatic activity (**SI Fig. 1a-c**). Single-molecule imaging of WT PBP1b revealed that the protein adopted a single major FRET state, centered at FRET efficiency ∼ 0.5 (FE, state 1), with sparsely populated tails at higher- and lower-FRET values (**Fig. 1c, SI Table 1**). Individual trajectories showed few – if any – transitions into and out of this state, and global analysis of transition frequency, depicted visually as a heatmap, showed that apo WT PBP1b was largely static (**Fig. 1d-e**). To rule out the possibility that PBP1 undergoes rapid dynamics undetected by our measurements, we imaged PBP1b at a 5-times faster frame rate (20 s^−1^) and confirmed that on its own the protein adopts a single stable conformation, i.e., an inactive or basal state (**SI Fig. 1d-e**).

**Figure 1:**
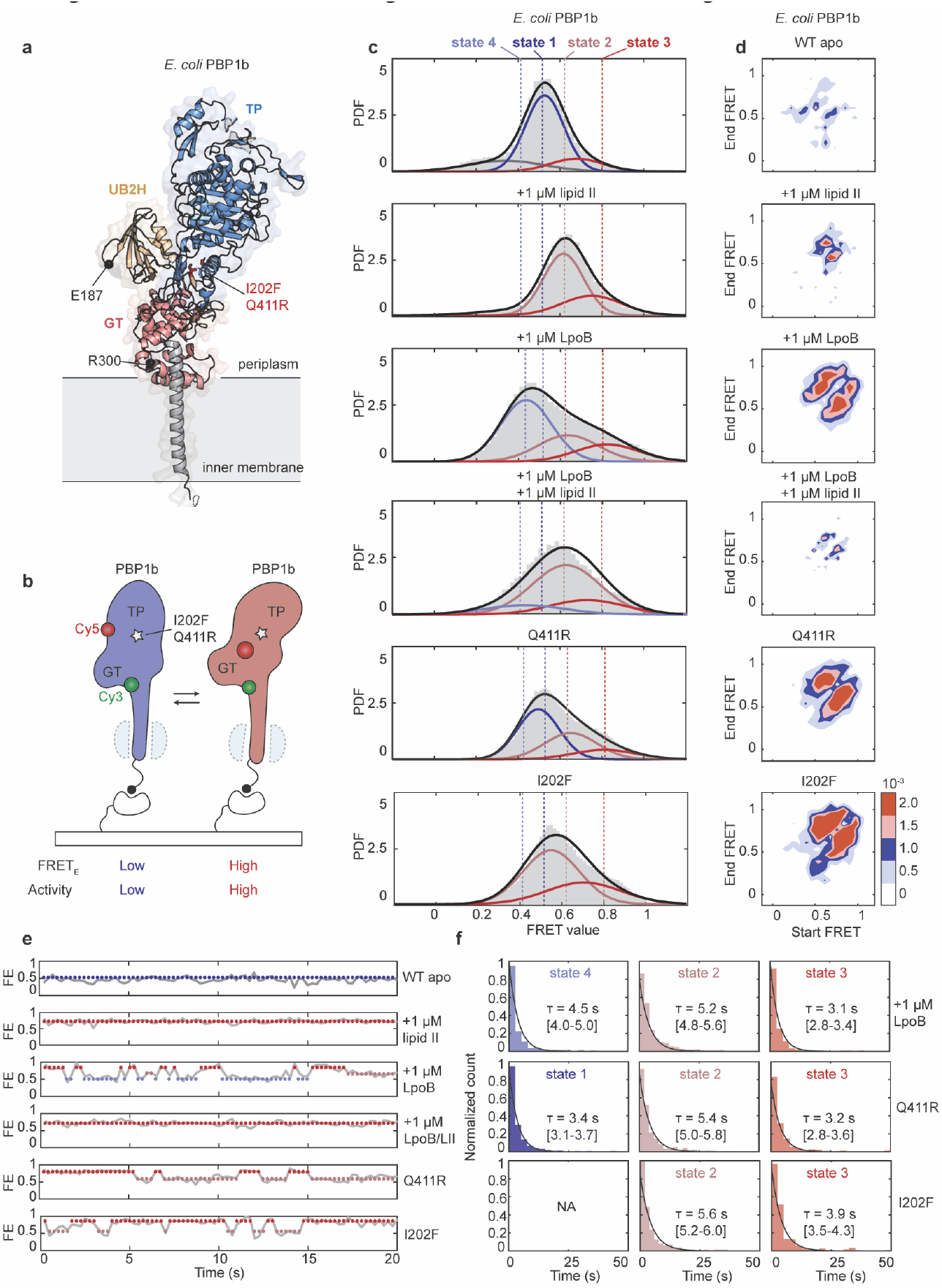
LpoB stabilizes the activated state of PBP1b. **a**, AlphaFold model of full-length *Ec*PBP1b showing the positions of cysteine substitutions (black spheres) and suppressor variants (red sticks). **b**, Schematic illustrating the smFRET assay, with one of the two possible orientations of donor (green) and acceptor (red) labels shown for simplicity. In the apo state, PBP1b adopts a lower-FRET efficiency state (inactive state). Activating perturbations (suppressor variants, lipid II or LpoB addition) shift the conformation of PBP1b to higher-FRET states. **c**, Probability density (PDF) histograms and fits of FRET efficiency (FE) values derived from single-molecule trajectories of *Ec*PBP1b^E187C-R300C^WT and suppressor variants (Q411R, I202F), with or without LpoB and lipid II. Normal fits to the data are shown in a blue-to-red color gradient, with states numbered 1-4. Mean values and occupancies of smFRET states are summarized in SI Table 1. **d**, Transition density heat maps, normalized to the total observation time, show the frequency of transitions for datasets in (**c**). White-to-red color gradient depicts the frequency of transitions from a starting FRET value (x-axis) to the final FRET value (y-axis), with white color corresponding to absence of transitions and red corresponding to high frequency of transitions. **e**, Representative single-molecule trajectories from datasets in (c). Markers are plotted at the mean values of state fits and colored as in (**c**). **f**, Dwell time histograms and fits for the states observed in datasets from (**c**). Mean dwell times alongside 95% confidence intervals are indicated on the plots. Conditions in which the protein remained largely static were omitted from analysis.

Next, we tested the effect of substrate addition on PBP1b dynamics by adding 1 µM lipid II into the flow cell. Lipid II binding resulted in a global structural shift to higher-FRET values, with a major state at FE ∼ 0.6 (state 2) and a minor state at FE ∼ 0.75 (state 3) (**Fig. 1c**). Both states exhibited few transitions, suggesting that lipid II binding stably captured a set of PBP1b conformations that were distinct from the apo state (**Fig. 1d-e**). Moreover, binding of a lipid II mimetic moenomycin, which was proposed to induce a reorientation of TP and GT domains^40,41^, led to a similar shift in the FRET ensemble (**SI Fig. 1d-e**). Thus, we conclude that a shift to higher-FRET values proceeds through the tilting of the TP domain closer to the GT domain and that this conformational change is a signature of enzymatic activation.

### LpoB binding to PBP1b promotes a structural transition to the activated state

LpoB is known to accelerate the GT activity of PBP1b *in vitro*^32,34-37^. We therefore aimed to assess whether addition of LpoB biases the conformation of PBP1b to an activated state. We found that LpoB binding partially enriched states 2 and 3, as well as a previously unobserved lower-FRET state (FE ∼ 0.4, state 4) (**Fig. 1c**). All three states exchanged rapidly on the timescales of our measurements, persisting for only a few seconds on average (**Fig. 1d-f)**. Simultaneous addition of LpoB and lipid II resulted in a distribution that was qualitatively similar to the lipid II-bound state, suggesting that state 4 is an on-pathway LpoB-bound intermediate that interconverts to the activated conformation (states 2 and 3) upon lipid II binding (**Fig. 1c-e**). Finally, single-molecule analysis of LpoB bypass variants^32^ (PBP1b^I202F^, PBP1b^Q411R^) showed that both substitutions dramatically shift the FRET distribution to the activated state and induce rapid dynamics within the protein (**Fig. 1d-f**), mimicking the activating effect of LpoB. Together, these results demonstrate that LpoB binding to PBP1b stabilizes the substrate-bound state of the enzyme through structural rearrangements at the GT-TP interface.

### Engineered variants of LpoB exhibit a range of affinities and activities

LpoB is an essential cellular activator of PBP1b and has been proposed to recruit PBP1b to the sites of cell wall damage^27,30,31,34^. Yet, LpoB binds PBP1b very weakly and is not strictly required for its GT activity *in vitro*^32,34–37^. We therefore sought to establish the extent to which recruitment and activation by LpoB contribute to the cellular function of PBP1b. For this purpose, we engineered LpoB variants with a range of recruitment and activation efficiencies, approximated by the ability of these variants to bind and induce conformational changes within PBP1b *in vitro*. Briefly, we designed a site saturation variant library of LpoB, targeting residues at the PBP1b interaction interfaces, displayed this library on yeast and selected high-affinity clones through two rounds of fluorescence-activated cell sorting (FACS)^43,44^ (**Fig. 2a, SI Fig. 2**, also *Methods*). We isolated 13 clones with substitutions at four positions (S157, A164, Y178 and S180) that mapped to an interface loop (termed loop 2) and a β hairpin (termed β3) of LpoB and proceeded with biochemical evaluation of the highest-affinity substitutions (**Fig. 2a-b**, also *Methods*). We also generated a non-binding LpoB control (E201R) by mutating the predicted hydrogen bond network at a PBP1b-LpoB interface (**Fig. 2a-b**, termed loop 4).

**Figure 2:**
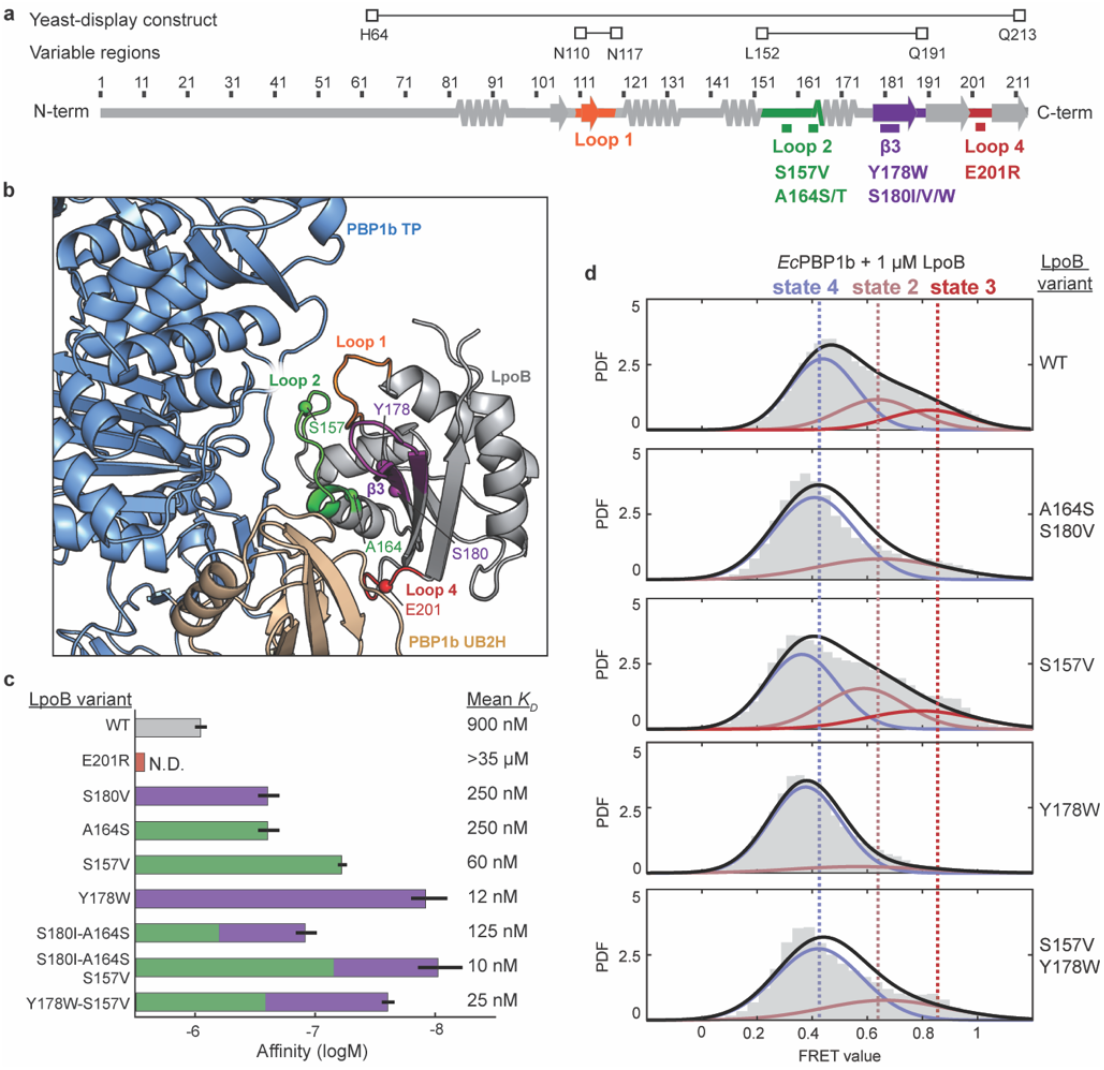
Affinity-matured LpoB variants stabilize active and inactive states of PBP1b. **a**, LpoB sequence, showing the truncation that was displayed on yeast for selections (LpoB^64-213^) as well as the variable regions of LpoB (LpoB^110-117^, LpoB^152-191^), in which all possible amino acid substitutions were incorporated at each position. Key interfaces are colored in orange (loop 1), green (loop 2), magenta (β3), and red (loop 4). **b**, AlphaFold model of the PBP1b-LpoB complex, showing the positions of the identified mutations as spheres. Interaction regions are colored as in (**a**). **c**, Bar graph summarizes BLI-determined affinities of LpoB variants color-coded by their positions in either loop 2 or β3, as in (**a**). Mean *K*_D_ values are indicated on the plot and full statistics can be found in SI Table 2. **d**, PDF histograms of FRET efficiency values derived from single-molecule trajectories of *Ec*PBP1b<SUP>E187C-R300C </SUP>in the presence of 1 μM LpoB WT or engineered LpoB variants. WT LpoB data are reproduced from Fig. 1 for convenience.

All engineered LpoB variants expressed well and displayed a monodispered peak on size-exclusion chromatography (**SI Fig. 3a-b**). We quantified binding affinities of individual variants and their combinations using a bulk bio-layer interferometry (BLI) assay (*Methods*). Variants with A164S and S180I/V substitutions modestly increased PBP1b affinity (K_*D*_ change from 900 nM to 250-300 nM), while the S157V and Y178W substitutions were more potent, improving affinity 15-75-fold (K_*D*_ = 12-60 nM), primarily by reducing the dissociation rate (**Fig. 2c, SI Fig. 3c, SI Table 2**). Combining the A164S, S180I, and S157 substitutions had an additive effect, yielding the highest affinity variant (K_*D*_ = 10 nM). In contrast, combining the S157V and Y178W changes was not additive (K_*D*_ = 25 nM), suggesting that the Y178W substitution does not stabilize the same PBP1b state(s) as the other three.

We next assessed how affinity-enhancing mutations affect the structural dynamics of PBP1b, and by extension, allosteric activation (**Fig. 2d**). We found that the four mutants varied in their ability to stabilize the activated state of PBP1b (FRET states 2 and 3) (**Fig. 2d, SI Table 1**). Specifically, the A164S/S180I double mutant partially stabilized the activated state; the S157V substitution was as effective as LpoB WT at activation, while the Y178W variant showed a near complete loss of the activated conformation. Combining the Y178W and S157V mutations partially restored the ability of the double mutant to stabilize the activated state of PBP1b. In summary, our selection campaign identified a set of functionally diverse LpoB variants. Notably, two mutants with significantly enhanced affinity for PBP1b had opposite effects on allosteric activation: Y178W showed an activation defect, whereas S157V fully retained activation.

### Decrease in LpoB-mediated activation reduces cell survival

We next aimed to correlate the biochemical properties of engineered LpoB variants with their activity *in vivo*, using a previously developed complementation assay^30^ (*Methods*). This assay takes advantage of the fact that *E. coli* cells remain viable if either PBP1a or PBP1b is inactive, but that inactivating both enzymes is lethal^28,29^. Thus, a *lpoB* deletion strain fails to grow when PBP1a (*ponA*) is depleted, but growth can be restored with expression of a functional copy of *lpoB*. In brief, we integrated^45^ genes encoding LpoB variants into the chromosome under the control of an inducible lactose promoter (P_lac_) in a PBP1a depletion strain deleted for *lpoB* (Δ*lpoB*Δ*ponA* P_ara_*::ponA*) and tested the ability of LpoB variants to complement growth in media and on plates (*Methods*). We found that growth phenotypes paralleled the ability of LpoB variants to activate, rather than to bind PBP1b. Affinity matured LpoB variants that maintained activation modestly improved growth, whereas the Y178W variant, expected to be highly efficient at PBP1b binding but defective in activation, showed a severe growth defect (**Fig. 3a**). This defect was reversed in the Y178W-S157V strain, where activation by LpoB is largely restored. Importantly, these phenotypes were not due to differences in protein expression, as all variants retained WT levels of expression (**SI Fig. 3d**). Together, these results establish that LpoB activation defects are detrimental to cell survival, whereas increased recruitment efficiency confers minimal advantages, suggesting that the low affinity of LpoB for PBP1b might be a beneficial feature of its regulation.

**Figure 3:**
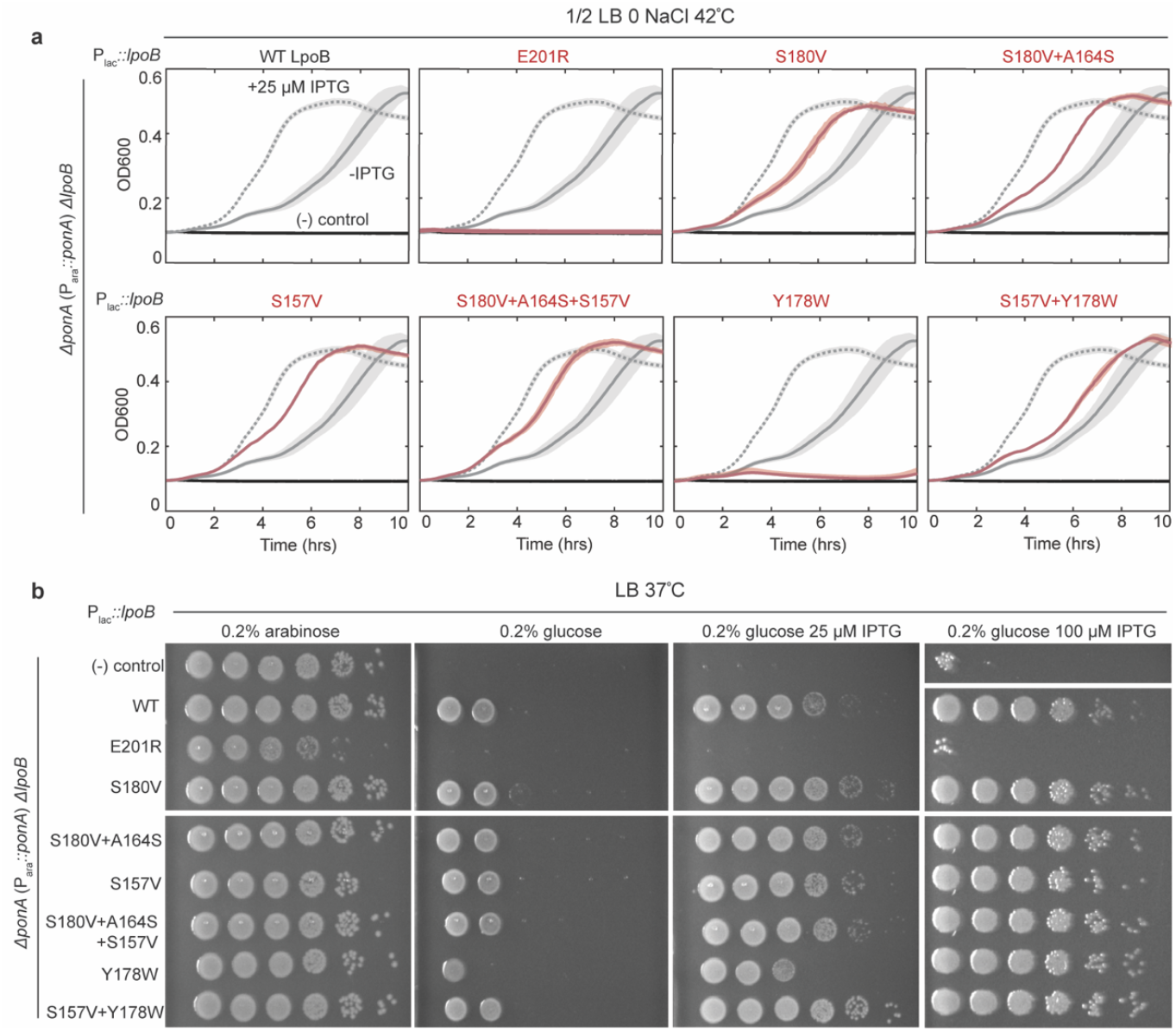
Cellular growth and survival scale with activation efficiency of LpoB variants. **a**, Growth curves under non-permissive growth conditions (½ LB 0 NaCl 42^°^C) of Δ*ponA* P_ara_*::ponA* Δ*lpoB* P_lac_::*lpoB* strains of negative control (black), LpoB WT (grey) or engineered LpoB variants (red). Averages of three technical replicates are shown as lines, with standard deviation depicted as a shaded region. In the absence of induction, strains of higher affinity variants that retain the ability to activate PBP1b exhibit modestly faster growth than the WT strains. Strains of LpoB mutants that disrupt binding to PBP1b (E201R) or enzymatic activation (Y178W) fail to grow. **b**, Titer experiments with strains from (**a**). Overnight cultures were serially diluted and spotted on LB agar with either 0.2% arabinose, 0.2% glucose, 25 μM IPTG or 100 μM IPTG. Data shown in (**a**) and (**b**) are representative of three biological replicates.

### PBP1b and LpoB form a transient complex

Given that LpoB-mediated activation is required for PBP1b catalysis *in vivo*, we sought to explore how LpoB contributes to each step of the polymerization reaction by measuring LpoB binding to PBP1b in the presence of lipid II substrate and glycan chain intermediates. To capture the dwell times of individual complexes, we developed a single-molecule interaction assay for PBP1b and LpoB (see *Methods*). To this end, PBP1b and LpoB were conjugated to Cy3- and Cy5-fluorophores, respectively, via cysteines engineered to produce a high-FRET signal upon complex formation. PBP1b-Cy3 was tethered to the flow cell surface, whereas LpoB-Cy5 was applied in solution, such that baseline Cy5- and FRET-signals remained near-zero and increased dramatically only upon LpoB binding (**Fig. 4a-b**). We confirmed that smFRET constructs were monodispersed by size-exclusion chromatography and fully retained binding and activity (**SI Fig. 4a-c**).

**Figure 4:**
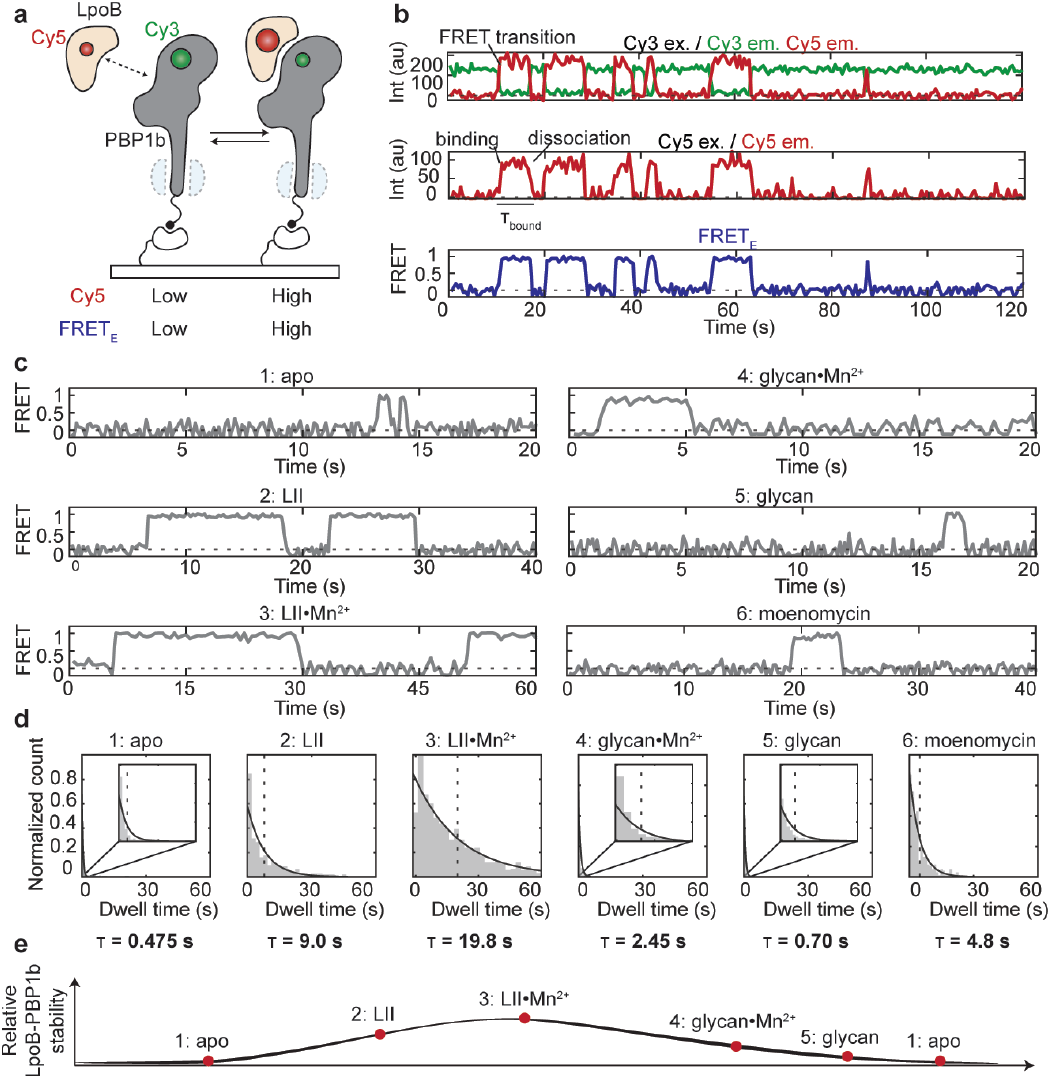
LpoB preferentially promotes glycan initiation over elongation. **a**, Schematic illustrating the PBP1b-LpoB smFRET binding assay. Cy3-labeled PBP1b is tethered to the surface, whereas Cy5-labeled LpoB is supplied in solution. In this set-up, Cy5 fluorescence and FRET signal are detected only when LpoB binds PBP1b. **b**, Example trajectories, showing donor and acceptor fluorescence upon donor excitation (top), accep-tor fluorescence upon acceptor excitation (middle) and the calculated FRET efficiency signal (bottom). Anti-correlated changes in donor and acceptor fluorescence (top) correspond to FRET transitions (bottom) and perfectly correlate with changes in direct acceptor signal (middle), i.e., Cy5-Cy3 colocalization events. **c**, Example trajectories of LpoB binding either to apo PBP1b or to PBP1b complexed with lipid II substrate or glycan chains. Step-like increases and decreases in FRET efficiency correspond to association and dissociation events, respectively. **d**, Dwell time histograms and exponential fits of samples from (**c**) with mean fits indicated on the figure. Insets show enlarged views of the histograms and fits in the 0 to 5 s range. Full binding statistics are summarized in the SI Table 3. **e**, Schematic illustrating the relative changes in the stability of the PBP1b-LpoB complex throughout the polymerization reaction.

Bulk BLI and smFRET assays yielded comparable *K*_D_ values for the WT PBP1b-LpoB interaction (*K*_D_ = 915 nM vs 900 nM) that closely match previously reported affinities^34,35^. However, we found that BLI consistently underestimated the dissociation rate of LpoB, likely due to avidity effects that arise from the high surface density of PBP1b required to achieve an adequate binding signal^46^. Our single-molecule assay, which is not prone to such artifacts, revealed that LpoB binds to PBP1b WT transiently, with binding events lasting on average less than half a second (**Fig. 4c-d, SI Table 3**). Bypass variants PBP1b^Q411R^ and PBP1b^I202F^ exhibit dramatically increased dwell times of the LpoB-bound state (τ = 7.7 and 10 s, respectively), consistent with the idea that LpoB has higher affinity for the activated state of the synthase^32,35^ (**SI Fig. 5**).

### LpoB preferentially promotes glycan initiation over elongation

With this assay in hand, we measured how the stability of the LpoB-PBP1b complex changes as the enzyme progresses through the polymerization reaction. We approximated the main stages of polymerization – substrate binding, initiation and elongation – by adding either lipid II substrate alone (binding), lipid II in the presence of divalent cations (initiation), or glycan chains linked to the lipid carrier (elongation). The addition of 1 µM lipid II increased the dwell time of the LpoB-bound state nearly 20-fold to τ = 9 s (**Fig. 4c-d**). This result demonstrates that LpoB has a higher affinity for the substrate-bound state of PBP1b and by extension, that LpoB binding promotes substrate recruitment to the enzyme. The substrate mimetic moenomycin also stabilized the complex between PBP1b and LpoB (**Fig. 4c-d**), suggesting that this antibiotic captures an on-pathway polymerization intermediate of PBP1b^42,47,48^.

Next, we added lipid II in the presence of manganese to allow polymerization to occur^49^. In our assay, lipid II is maintained in the flow cell in overwhelming stoichiometric excess over the enzyme (∼20,000x), ensuring that any short chains that are formed are rapidly exchanged for monomeric lipid II and allowing us capture the early stages of the polymerization reaction. We observed a further increase in the dwell time of the LpoB-PBP1b complex to τ ∼ 20 s (**Fig. 4c-d**), indicating that LpoB promotes synthesis initiation in addition to substrate binding.

Finally, to approximate the LpoB-PBP1b interaction during the elongation stage of polymerization, we supplied lipid-linked glycan chains (10-15 headgroups in length) into the flow cell, matching the concentration of the lipid carrier to that of the free lipid II used above (**SI Fig. 4d**, also *Methods*). We found that the dwell times of the glycan-bound states both with (τ = 2.5 s) and without (τ = 0.7 s) manganese were considerably shorter than those of lipid II-bound state (**Fig. 4c-d**). In agreement with this result, addition of glycan chains did not stabilize the activated conformation of PBP1b in smFRET dynamics assays (**SI Fig. 1d-e**). We conclude that LpoB preferentially associates with the substrate-bound state of the enzyme and promotes glycan synthesis initiation, but likely does not contribute significantly to elongation (**Fig. 4e**). This result explains earlier observations in bulk polymerization assays that addition of LpoB shifts the distribution of glycan chains to shorter length, presumably due to an increase in the rate of initiation (**SI Fig. 4c**)^32^.

### Synthesis time by PBP1b is independent of its affinity for LpoB

Our smFRET experiments strongly suggest that LpoB functions primarily during the early stages of PG synthesis, but dissociates from PBP1b at the later stages of PG synthesis. To investigate how changing LpoB affinity affects PG synthesis in cells, we performed single-particle tracking (sp-tracking) of PBP1b WT and the activating I202F variant which increases the dwell time of the LpoB bound state 20-fold (**SI Fig. 5b**). To that end, we introduced a functional Halo-tagged PBP1b fusion and its I202F variant into the background of a *ponB* deletion strain and imaged the resulting strains with a fast acquisition frame rate (30 ms) to quantify the fractions of mobile and immobile molecules (**SI Fig. 6a-b, SI Table 4**, also *Methods*). Earlier sp-tracking studies established that immobile (bound) foci of PBP1b correspond to either LpoB-tethered or actively-synthesizing enzymes, whereas highly diffusive particles are inactive^21,27,50^. We observed a modest, but reproducible increase (∼15-20%) in the fraction of the bound state for the I202F variant in comparison to WT PBP1b, consistent with an increase in synthesis initiation conferred by the mutation (**Fig. 5a**). We note that these population differences were not due to changes in expression levels, as I202F and WT exhibited equivalent levels of expression across a range of induction conditions (**SI Fig. 6e**).

**Figure 5:**
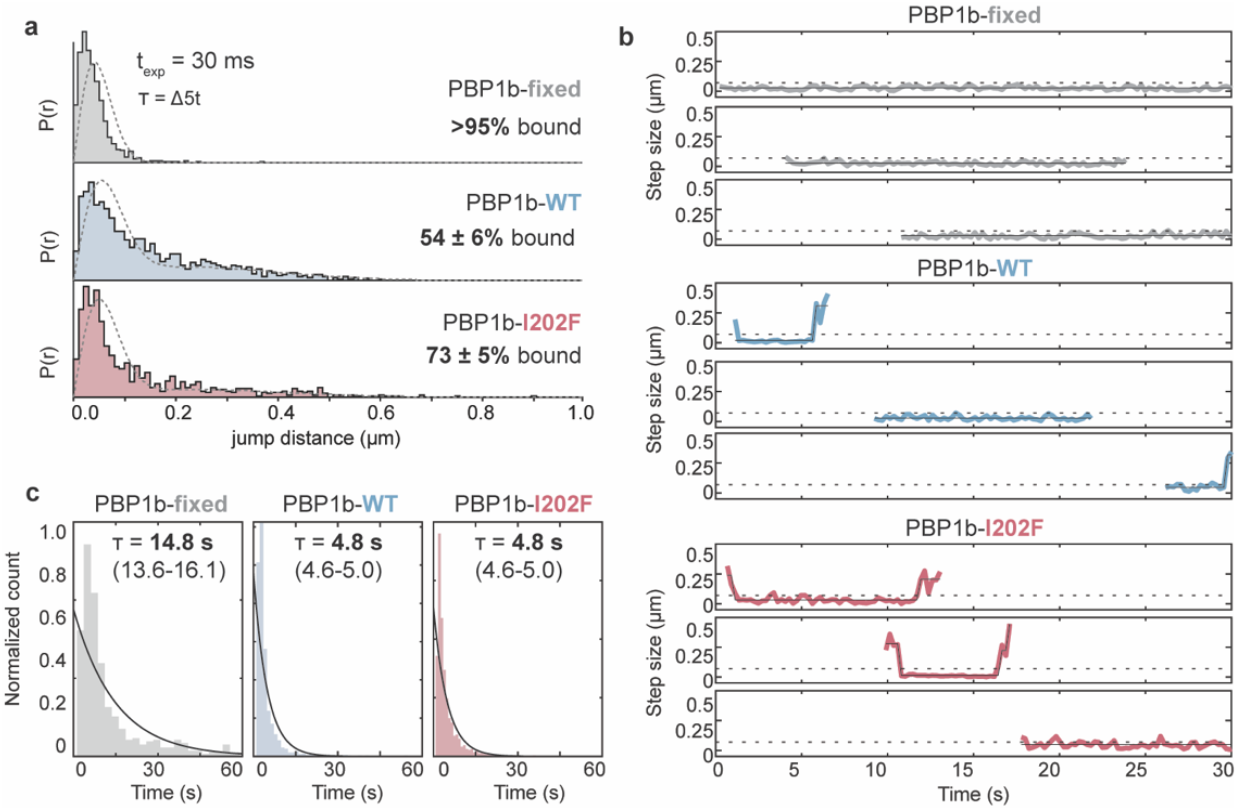
PBP1b synthesis time is independent of its intrinsic affinity for LpoB. **a**, Representative jump distance distributions and fits of strains expressing PBP1b-Halo-WT (TU122/pAV23, blue) and PBP1b-Halo-I202F mutant (TU122/pSI237, red), alongside the fixed cell control (TU122/pAV23, grey), imaged with a fast acquisition rate (30 ms). The I202F strain exhibits a 15-20%-fold increase (p < 0.05) in the population of the bound (active) state relative to the WT sample. Data collection parameters and statistics are summarized in SI Table 4. Distributions and fits were generated using Spot-ON sp-tracking software. **b**, Example single-particle trajectories of strains from (**a**) of PBP1b-Halo-WT (blue), PBP1b-Halo-I202F mutant (red) and the fixed cell control (grey) imaged with a slower acquisition rate (225 ms). Periods of immobility (active synthesis or tethering to LpoB) correspond to step-like decreases in the instantaneous step size of the diffusing particle below the threshold value of 70 nm (dotted line). The fixed cell control shows few if any rapidly diffusing particles and has much longer periods of immobility. **c**, Dwell time distribution histograms and exponential fits of samples from (**b**). PBP1b-WT and the I202F mutant exhibit similar dwell times of the bound (active) state.

We next asked whether increasing PBP1b affinity for LpoB leads to longer dwell times of the immobile PBP1b population due to either extended synthesis or LpoB-trapping. To measure the immobile state dwell times, we imaged the WT and I202F strains with longer exposure times (225 ms) and generated trajectories of step size as a function of time for each particle (**SI Fig. 6b**, see *Methods*). These trajectories showed reversible transitions between highly diffusive and immobile states occurring on the timescales of seconds, with drops in step size corresponding to periods of immobility (**Fig. 5b**). We defined an ‘immobilization’ threshold (70 nm) based on the jump distance distribution of a paraform-aldehyde-fixed control, and quantified the dwell times of events that fell below this threshold and lasted at least four frames (**SI Fig. 6c-d**, also *Methods*). Notably, PBP1b WT and the I202F variant exhibited equally transient immobile states that persisted for only 5 seconds on average, indicating that increasing LpoB affinity did not lengthen total synthesis time or cause PBP1b to be trapped in complex with LpoB (**Fig. 5c, SI Table 4**). The observed dwell times were not limited by photobleaching, since particles in the fixed control sample were found to persist for much longer periods of time (**Fig. 5b-c**). Collectively with our *in vitro* data, these results suggest that LpoB transiently binds PBP1b promoting initiation but does not remain associated with the enzyme throughout the entire process of synthesis.

## Discussion

The prototypical *E. coli* synthase PBP1b has been extensively studied genetically and biochemically, yet the molecular mechanism by which its lipoprotein activator LpoB stimulates PG repair in cells has remained unclear. Here we show that LpoB functions as an initiator of PG synthesis, promoting site-specific recruitment and enzymatic activation of PBP1b through structural rearrangements at the GT-TP interface.

The low affinity of apo PBP1b for both its substrate and LpoB is likely a central and necessary feature of its regulation, ensuring that untargeted PG synthesis is minimized (**Fig. 6**, search and recruitment). *In vitro*, however, PBP1b binds lipid II and transitions to the activated state in the absence of LpoB, resulting in non-negligible polymerization activity (**Fig. 1, SI Fig. 1**). The discrepancy between *in vitro* and *in vivo* behavior can be accounted for by differences in substrate concentration between the two environments. In *E. coli*, lipid II is present at a copy number of ∼1000-2000 molecules per cell, similar to the total number of PG synthases that compete for it and 3-4 orders of magnitude lower than what was used in our smFRET experiments or in detergent-based GT assays^32,35–37,51–53^. Thus, in the absence of membrane partitioning that would elevate local substrate concentration, binding events between lipid II and the apo enzyme in cells are likely too infrequent and short-lived to permit simultaneous recruitment of two lipid II molecules required to initiate synthesis^54^. At sites of synthesis, LpoB binding induces structural changes in PBP1b that increase its affinity for substrate 20-40-fold and dramatically enhance initiation efficiency (**Fig. 6**, initiation). As the polymerization reaction progresses, the stability of the LpoB-PBP1b complex decreases, and LpoB dissociates from the synthesizing enzyme (**Fig. 6**, LpoB recycling). Rapid dissociation ensures that LpoB does not become trapped in the PG matrix and is efficiently redistributed to other areas in need of fortification.

What allows the later stages of synthesis to continue without LpoB? We show that individual bursts of synthesis last only a few seconds and likely do not involve processive polymerization. Indeed, AFM studies estimate that most defects in the *E. coli* PG matrix are ∼10-15 nm^2^ in size, requiring incorporation of glycan material only a few lipid II headgroups long^55^. We therefore propose that after LpoB dissociates, tethering to the nascent glycan chain and the PG meshwork retains PBP1b at the site of synthesis long enough to complete repair (**Fig. 6**, synthesis completion). Such a mechanism, in which the synthesis activity of PBP1b is not limited by the intrinsic dwell time of the LpoB-bound state, allows the enzyme to patch defects of varying sizes. Future work will determine how additional modulators of PBP1b activity, e.g., CpoB, may interplay with LpoB-mediated regulation of activity and spatiotemporal distribution of aPBPs^35,56^.

**Figure 6:**
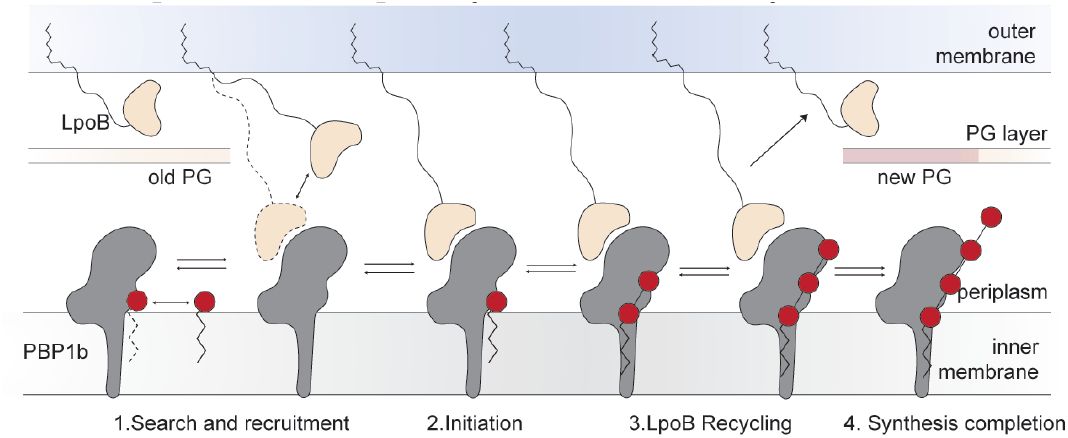
LpoB forms a transient complex with PBP1b, triggering synthesis initiation. Schematic overview shows the model of LpoB-mediated PG synthesis by PBP1b. Outside of areas of low PG density, low substrate availability and low PBP1b affinity for lipid II prevent the enzyme from efficiently initiating synthesis. Once PBP1b encounters LpoB at a gap in the PG matrix, LpoB binding allosterically increases substrate affinity and promotes initiation. As polymerization progresses, LpoB affinity for PBP1b decreases and it is recycled from the synthesizing complex before PG synthesis can close off its escape route. PBP1b is retained on the PG substrate and completes synthesis.

Our smFRET experiments demonstrate that PBP1b assumes a range of structural states with distinct functional roles. The higher FRET states, in which GT and TP-UB2H domains are brought closer together, correspond to the activated conformations of the enzyme, whereas the lower FRET states are inactive. This finding is in broad agreement with previously determined apo and moenomycin-bound structures of *Ec*PBP1b that show that antibiotic binding is accompanied by a rigid-body tilting of TP and UB2H domains towards the GT domain^40–42^. In addition to these two states, we detected a previously unobserved conformation that PBP1b adopts upon binding to LpoB. The LpoB variant (Y178W) that captures this state has a loss-of-function phenotype in cellular assays, suggesting that this conformation corresponds to an inactive on-pathway intermediate that transitions to the activated state in the presence of lipid II. Future structural studies will be required to determine how LpoB-induced conformational changes are propagated to the active site of PBP1b to promote initiation, as well as how this coupling is lost at the later stages of synthesis. Due to the transient nature of their interaction, PBP1b complexes with LpoB and substrate have eluded structure determination. Our high-affinity LpoB variants provide an invaluable structural tool for stabilizing distinct functional intermediates of PBP1b to enable atomic-level resolution of the PG synthesis process. Finally, engineered LpoB variants that trap active and inactive conformations of PBP1b can be leveraged in high-throughput small molecule screens to identify novel inhibitors of PBP1b.

Precise spatiotemporal coupling of GT and TP reactions to accessory factor binding is critical for proper PG assembly^57^. We propose that structural dynamics at the GT-TP interface represent a unifying regulatory mechanism that coordinates activation of polymerization and crosslinking reactions by different classes of PG synthases, responsible for elongation, division, and PG fortification and repair. Indeed, activating mutations that map to this interface have been identified in several aPBPs as well as the structurally unrelated SEDS-bPBP enzymatic complexes from both gram-negative and gram-positive bacteria (**SI Fig. 7**)^19,32,33,39,58–62^. In each system, the allosteric switch is adapted to fit the specific mechanistic requirements of the enzyme’s physiological function. For instance, unlike PBP1b, SEDS-bPBPs undergo large-scale conformational rearrangements to achieve enzymatic activation and form stable complexes with their cellular partners to carry out processive PG synthesis^21,24,25,38,63^. We speculate that the conservation of this regulatory mechanism across a diverse range of structural folds and functions arises from the shared requirement for concerted activation of polymerization and crosslinking reactions.

## Acknowledgements

Single-particle tracking data were collected at the Microscopy Resources on the North Quad (MicRoN) core at Harvard Medical School. BLI experiments were performed in the Center for Macromolecular Interactions Core Facility (RRID:SCR_018270) at Harvard Medical School. I.S. is supported by the Van Maanen Fellowship from the Department of Biological Chemistry and Molecular Pharmacology at Harvard Medical School and the National Science Foundation (NSF) Graduate Research Fellowship award. A.V. was supported in part by a EMBO long-term postdoctoral fellowship (ALTF_89-2019) and a SNF Postdoc Mobility fellowship (P500PB_203143). Funding for this work was provided by National Institutes of Health grants U19 AI158028 (A.C.K. and T.G.B.), R01 GM114065 (J.J.L.), R01 AI083365 (T.G.B.), and investigator funds from the Howard Hughes Medical Institute (T.G.B.). We thank members of the Bernhardt, Loparo, and Kruse labs for their helpful suggestions and members of the MicroN and CMI facilities for their support. This paper was typeset with the bioRxiv word template by @Chrelli: www.github.com/chrelli/bioRxiv-word-template.

## Competing interests

A.C.K. is a cofounder and consultant for biotechnology companies Tectonic Therapeutic and Seismic Therapeutic, and for the Institute for Protein Innovation, a non-profit research institute.

## Data availability

Custom single-particle tracking and single-molecule FRET Matlab scripts are available upon reasonable request.

## Materials and Methods

### Construct design for smFRET experiments

All constructs used for smFRET microscopy were assembled via PCR and Gibson assembly (SI Table 5). Previously described soluble truncations of *Ec*PBP1b^58-804^ and *Ec*LpoB^21-213^ (pCB39) were used as background constructs for all *in vitro* applications, unless otherwise specified^30,40-42^. For smFRET dynamics measurements, a structural disulfide in PBP1b was substituted with a salt bridge (*Ec*PBP1b^C776R-C794D^) to prevent nonspecific labeling. Two cysteines were introduced at positions that do not interfere with enzymatic activity or LpoB binding to monitor conformational changes within PBP1b (*Ec*PBP1b^E187C-R300C^). Finally, N-terminal FLAG and ALFA tags were incorporated into this construct (pSI228) for M1 affinity purification and immobilization on strep-tavidin-functionalized coverslips via a biotinylated α-ALFA nanobody (NanoTag)^64^. Suppressor variants, I202F (pSI229) and Q411R (pSI265) were introduced into the background of the pSI228 “WT” variant. For smFRET binding experiments, single-cysteine substitutions in LpoB (pSI204, *Ec*LpoB^S211C^) and PBP1b (pSI233, *Ec*PBP1b^C794A^) were designed, based on the PBP1b-LpoB AlphaFold prediction, such that the distance between fluorophores would be 30 Å or less in the complex. For *Ec*PBP1b, one of the cysteines (C776) from the native disulfides was used for labeling, whereas the other cysteine (C794) was mutated to alanine. Suppressor variants, I202F (pSI234) and Q411R (pSI242) were introduced into the background of the pSI233 “WT” variant.

### Expression and purification of *Ec*PBP1b and *Ec*LpoB constructs

Expression and purification of constructs used for biochemistry and single-molecule imaging was carried as described before^19,38,65^. In brief, plasmids encoding *E. coli* PBP1b or LpoB variants (**SI Table 5**) were transformed into *E. coli* C43 (DE3) cells with or without SUMO tag-specific Ulp1 protease under an arabinose inducible plasmid (pAM174) respectively and grown at 37 °C for ∼16 h on LB-agar plates supplemented with appropriate antibiotics (100 µg/ml ampicillin, 35 µg/ml chloramphenicol, and 50 µg/ml kanamycin). In all subsequent steps, antibiotics were added to the media at the indicated dilutions. Transformants were scraped off the plate, inoculated into 5 ml LB pre-cultures and grown for 1 h at 37 °C with shaking. These cultures were diluted into 1 L of TB 2 mM MgCl_2_, 0.1% glucose, and 0.4% glycerol, and grown at 37 °C with shaking until an OD_600_ > 2. After the cells were cooled down to 18 °C, protein expression was induced by adding 1 mM isopropyl β-d-1-thiogalactopyranoside (IPTG) and/or 0.1% arabinose to the culture. After ∼16 h of induction at 18°C, bacterial pellets were harvested by centrifugation at 7,000 x g for 15 min and stored at −80 °C for subsequent purification.

For antibody-based affinity purification, bacterial pellets were resuspended in 50 mM HEPES pH 7.5, 150 mM NaCl, 20 mM MgCl_2_ (lysis) buffer supplemented with protease inhibitors tablets (ThermoFisher) and benzonase nuclease (∼1.5 units/ml, Sigma Aldrich). For, nickel-based purification, 50 mM Tris pH 7.5, 300 mM NaCl lysis buffer was used instead. Resuspended bacteria were homogenized and lysed by three passes through an LM10 microfluidizer (Microfluidics) at 15,000 psi. The lysate was then centrifuged for 1 h at 50,000 x g to separate membrane (pellet) and cytoplasmic (supernatant) cellular fractions. For soluble constructs (LpoB variants), the supernatant was collected, passed through a fiberglass filter and supplemented with 5 mM imidazole (nickel) or 2 mM CaCl_2_ (anti-Protein C) prior to affinity purification. For membrane protein constructs (PBP1b variants), the supernatant was discarded and membrane pellets were resuspended in 20 mM HEPES pH 7.5, 0.5 M NaCl, 5% glycerol, 1% n-dodecyl-β-D-maltopyranoside (DDM) buffer with douncing (ThermoFisher). The resulting mixture was solubilized for ∼2 h at 4°C with stirring and centrifuged at 50,000 x g for 1 h to separate soluble (supernatant) and insoluble fractions. The supernatant was filtered and supplemented with 2 mM CaCl_2_ prior to loading onto affinity resin.

For nickel-based purification, supernatant was loaded onto 1-2 mL of nickel resin previously equilibrated with lysis buffer supplemented with 5 mM imidazole. The column was then washed with 10-20 column volumes (CV) of equilibration buffer, and protein was eluted with 200 mM imidazole in 20 mM Hepes pH 7.5, 300 mM NaCl buffer. For antibody-based purifications, supernatants were loaded onto 2-4 mL of anti-protein C tag or M1 antibody resins equilibrated with 20 mM Hepes pH 7.5, 300 mM NaCl buffer with (membrane constructs) or without (soluble constructs) 0.1% DDM supplemented with 2 mM CaCl_2_. The columns were washed with 10-20 CVs of equilibration buffer, and proteins were eluted with 10 CVs of 0.2 mg/mL FLAG peptide or 0.2 mg/mL Protein C peptide in 20 mM Hepes pH 7.5, 300 mM NaCl 5 mM EDTA with or without 0.1% DDM. Eluates were concentrated to ∼ 500 µl using an Amicon Ultra Centrifugal filter (Millipore Sigma) with a 10 kDa molecular weight cutoff (MWCO) for LpoB samples and a 100 kDa MWCO for PBP1b samples. All samples were additionally purified on a Superdex 200 Increase (Cytiva) SEC column using 20 mM HEPES pH 7.5, 350 mM NaCl buffer with or without 0.02% DDM; monomeric peak fractions were concentrated as before and flash frozen for later use.

### GT activity assays

Lipid II fluorescent labeling and GT assays were performed as described previously^38^. Briefly, 10 µM stocks of SEC-purified *Ec*PBP1b (see above) were diluted 10-fold into 20 mM Hepes pH 7.5, 300 mM NaCl 0.02% DDM buffer containing 10 µM lipid II-AF488 and 2 mM MnCl_2_ and either 5 µM LpoB or buffer control and incubated for 10-30 min at 25 °C. The reactions were quenched by incubation at 4 °C and the addition of SDS loading dye. The samples were then loaded into a 4–20% gradient polyacrylamide gel and run at 200 V for 30 min, after which glycan chains were visualized using the fluorescence Typhoon imager (Amersham Typhoon 5).

### Glycan chain preparation

100 µL of 200 µM lipid II in DMSO was diluted 1:5 into 0.02% DDM buffer and polymerized by 20 µM PBP1b for 30 minutes as above. Glycan products were loaded onto a C18 BakerBond column (VWR) equilibrated with 5 CV of MeOH/0.1% NH_4_OH, followed by 5 CV of H_2_O/0.1% NH_4_OH. The column was washed with 2 CV of H_2_O/0.1% NH_4_OH, and glycan chains were separated from monomeric lipid II with a methanol gradient elution (0, 20, 40, 60, 80, 100 % MeOH/0.1% NH_4_OH). Two 1 CV fractions were collected for each methanol concentration, dried on a microvap for 2-6 h, lyophilized overnight, resuspended in 10 µL DMSO and stored at −20°C. Biotinylation and visualization of the resulting products were carried out following a previously described protocol^19^. Briefly, 1 µL of each fraction was diluted 10-fold into 20 mM Hepes pH 7.5, 300 mM NaCl buffer to which biotinylated d-lysine and *E. faecalis* PBPX were added at 2 mM and 10 µM working concentration, respectively. Following incubation for 30 min at 25°C, the reaction was quenched by addition of SDS loading dye, and samples were separated on SDS-PAGE. The products were transferred onto a low-fluorescence PVDF membrane (BioRad) and blocked in 3% BSA 1xPBS for 1 h at room temperature. Biotinylated products were visualized by incubation with fluorescently tagged streptavidin (1:5,000 IRDye 800-CW streptavidin, Li-Cor Biosciences) for an additional 1 h at room temperature. Membranes were washed three times with 1xPBS and imaged on the Typhoon imager.

### smFRET sample preparation and labeling

Constructs that required fluorophore labeling were purified as described above, up until the step of concentrating the eluates from M1 or anti-Protein C tag antibody columns. These eluates were incubated with 5 mM DTT for 30 minutes at room temperature (RT) to reduce any oxidized cysteines and purified on SEC using 20 mM HEPES pH 7.5, 350 mM NaCl 5 mM EDTA buffer with or without 0.02% DDM (SEC buffer). Main peak fractions were collected, concentrated and labeled with 10-fold stoichiometric excess of sulfo-Cy3-maleimide and/or sulfo-Cy5-maleimide fluorophores (Lumiprobe) for 15 min at RT. Unreacted dyes were removed using zeba spin desalting columns (ThermoFisher) and the samples were further purified on SEC. Monomeric peak fractions were concentrated to 10 µM and flash frozen in aliquots for smFRET experiments.

### smFRET chamber preparation and data collection

Microfluidic chambers for smFRET imaging were built as previously described^38,65^. Prior to imaging, the chamber was blocked with 1 mg/ml bovine serum albumin solution (NEB) for 5 min, followed by two washes with SEC buffer. Next, the surface was functionalized with 0.25 mg/ml streptavidin for 5 min, followed by two washes with SEC Buffer to remove unbound streptavidin.

Next, 0.1 mg/ml anti-ALFA tag biotinylated nanobody was added, incubated for 5 mins, and excess nanobody was washed off with SEC buffer. Following this, labeled PBP1b was added to the chamber at 100-500 pM concentration and incubated with a reactive oxygen species scavenging cocktail consisting of SEC buffer with 5 mM protocatechuic acid (PCA), 0.1 µM protocatechuate-3,4-dioxygenase (PCD), 1 mM ascorbic acid (AA), and 1 mM methyl viologen (MV) for 5 min. Images were collected on an Olympus IX-71 total internal reflection (TIRF) microscope using Hamamatsu HCImage live version 4.4.0.1 and Labview version 15.0f2 software as previously described^38,66^.

For smFRET dynamics measurements, the power was set to 2 W/cm^2^ for the 532 nm laser and 1 W/cm^2^ for the 641 nm laser. For each sample, movies of 60-300 seconds in length were collected at a frame rate of 4 s^−1^ with two frames of 532 nm excitation alternating with one frame 641 nm excitation. Fast imaging of the PBP1b WT sample was carried out with a frame rate of 20 s^−1^ and with 2-fold higher laser powers (4 W/cm^2^ for 532 nm laser and 2 W/cm^2^ for 641 nm laser).

For smFRET binding measurements, movies of 20-120 seconds in length were collected with alternating single frames of 532 and 641 excitation and at varying frame rates (4-20 s^−1^), adjusted to capture individual binding events for each condition **(SI Table 3**). PBP1b WT (pSI233) sample was additionally imaged at a frame rate of 20 s^−1^ with continuous 532 nm excitation to achieve maximal resolution of the transient binding events (τ_mean_ < 0.5 s). In those experiments, binding events were detected through FRET signal only. Laser powers were adjusted depending on the chosen exposure time. 2 W/cm^2^ (532 nm) and 1 W/cm^2^ (641) powers were used for 4 s^−1^; 3 W/cm^2^ (532 nm) and 1.5 W/cm^2^ (641) powers were used for 10 s^−1^; 4 W/cm^2^ (532 nm) and 2 W/cm^2^ (641) powers were used for 20 s^−1^.

### Analysis of smFRET dynamics experiments

Previously described Matlab analysis pipeline was used to perform spot detection, channel alignment and background subtraction of movies from **Fig. 1** and **Fig. 2**^38,66^. The resulting single-molecule trajectories were further filtered to exclude trajectories containing more than one donor and one acceptor fluorophore and cropped at the time of fluorophore photobleaching using a custom Matlab script^65^. The publicly available ebFRET GUI was used to analyze each dataset to determine the underlying distributions of smFRET states and the kinetics of their exchange in an unbiased manner^67^. Dwell-time histograms and exponential fits were generated by selecting time segments lasting at least three frames from trajectories exhibiting at least one true transition event. Transition frequency heat plots were generated by quantifying the total number of transition events observed in a dataset and normalizing by the total observation time. We note that dwell time histograms were not reported for *Ec*PBP1b “WT” apo and lipid II-containing samples, since in those conditions the transition frequency was so low that the dwell times were limited by fluorophore photobleaching. smFRET dynamics fits and statistics are summarized in **SI Table 1**.

### Analysis of smFRET binding experiments

Movies from **Fig. 4** were analyzed using a publicly available automated pipeline to calculate background-subtracted intensities of donor (PBP1b) and acceptor (LpoB) fluorophores as a function of time^68^. Resulting trajectories were further corrected for bleed through from donor channel to acceptor channel, direct excitation of acceptor fluorophore by donor excitation laser, and differences in fluorophore quantum yield and detection efficiency^69^, using a custom Matlab script. PBP1b-LpoB binding events were detected from donor and acceptor colocalization events and the concomitant FRET transitions. To exclude nonspecific colocalization events, thresholds were set for minimum acceptor fluorescence intensity, for the minimum FRET signal, and for the maximum displacement between acceptor and donor centroids. Dwell times histograms and exponential fits of the bound states were calculated from time segments lasting at least three consecutive frames (**SI Table 3**).

### LpoB affinity maturation

A 898-member site saturation variant library (SSVL) library in which all possible amino acid substitutions were incorporated into loops 1 and 2 and the beta hairpin of *Ec*LpoB^64-213^ was synthesized as double-stranded DNA (Twist Bioscience). Background vector for yeast-display (pYDS)^43^ was amplified and co-transformed with the insert DNA library into BJ5465 yeast with ECM 830 Electroporator (BTX-Harvard Apparatus). Yeast cells were recovered in YPAD for 1.5 h. Transformed cells containing the insert were selected by plating on tryptophan-deficient media.

Selections of high-affinity LpoB variants were carried out similarly to the nanobody campaigns described previously^44,70^. In brief, 10^8^ yeast cells from the LpoB-display library were harvested and resuspended in selection buffer (20 mM HEPES pH 7.5, 300 mM NaCl, 2.8 mM CaCl_2_, 0.1% DDM, 0.1% BSA, 0.2% maltose) and stained for 30 min at 4°C with either 100 nM or 25 nM of ALFA-tagged *Ec*PBP1b-WT labeled with streptavidin-AF488 via a biotinylated α-ALFA nanobody (**SI Fig. 2a**). Yeast cells were washed three times with selection buffer to remove unbound antigen and sorted on Sony SH800 cell sorter using SH800 software v.2.1.6 to select clones with increased binding affinity for PBP1b (**SI Fig. 2b**). Sorted cells were amplified and bulk titrations on the two FACS rounds, together with the naïve library control, were carried to assess population changes in affinity (**SI Fig. 2e**). Since both FACS rounds showed improvement in binding affinity for PBP1b, single-yeast staining experiments of 48 randomly selected clones from the two rounds of selections were used to isolate top binders (**SI Fig. 2c-d**). A threshold on the total bound fraction (2-fold higher than the naïve library) was set to exclude variants with negligible changes in binding affinity, and 13 clones that passed this filtering criterion were sequenced. All 13 contained substitutions at 4 positions (A164, S157, Y178, S180). Variants at the four positions (A164S, S157V, Y178W, S180I/V) with the highest on-yeast binding were cloned into bacterial expression plasmids for further biochemical analysis.

### Bio-layer interferometry (BLI) binding assays

The interaction affinity between PBP1b and LpoB variants was measured on Octet Red384 using the Octet BLI Discovery Software. PBP1b constructs, in which the TM domain was substituted with an N-terminal cysteine acylation site (*Ec*PBP1b^82-804^, pSI199), were used for these measurements to reduce nonspecific sticking of the membrane protein to the tip surface. Engineered variants of LpoB were produced in the background of a truncation used for affinity maturation (LpoB^64-213^, pSI259) to match yeast selection conditions. We confirmed that lipidated PBP1b and truncated LpoB bound with a similar affinity (900 nM) to what was previously reported for WT proteins^35^. BLI experiments were carried out in 20 mM Hepes pH 7.5 300 mM NaCl 0.1% DDM buffer (BLI buffer). First, 5 µg/mL anti-ALFA tag biotinylated nanobody in BLI buffer was immobilized on the surface of streptavidin-coated biosensors (Sartorius) to produce a total response of 1-2 nm. Following a wash to remove uncomplexed nanobody, 5 µg/mL ALFA-tagged PBP1b in BLI buffer was applied to the surface (4 nm total response). Following another wash to remove excess PBP1b, 8-point LpoB titrations in BLI buffer were carried with concentrations ranging from 3 nM to 5 µM, depending on condition. Resulting curves were exported to Octet Analysis Studio 13.0 Software, double-reference subtracted and fit to a global-fit model to extract kinetic and thermodynamic parameters of binding (**SI Table 2**).

### Preparation of *in vivo* strains

Plasmids and strains used for *in vivo* experiments can be found in **SI Tables 6** and **7**. For sp-tracking experiments, a sandwich Halo-Tag fusion of the *Ec*PBP1b gamma isoform (pAV23) and its I202F variant (pSI237) were CRIM-integrated^45^ into the background of the *ponB* deletion (TU122). Briefly, the host strain (TU122) was transformed with a helper plasmid (pAH69)^45^, and ampicillin-resistant transformants were selected on LB-agar with 100 µg/mL amp at 30°C. The resulting strain was used for CRIM-integration of target plasmids (pAV23, TU122) at the HK022 site. Following transformation, cultures were incubated overnight on LB-agar with 35 µg/mL chloramphenicol at 37°C to cure the helper plasmid. Cured derivatives (TU122/pAV23, TU122/ pSI237) were verified as antibiotic sensitive and streak purified. Single integration was confirmed by colony PCR, and expression was verified through western blotting.

Strains for LpoB complementation titers and growth curve assays were prepared as above, through CRIM-integration of LpoB variants under the control of the IPTG-inducible lac promoter (Plac) into the background of a *lpoB* deletion and *ponA* depletion strain (CB4)^30^ and selection with 10 µg/mL tetracycline (tet^10^) on LB-agar supplemented with 0.2% arabinose. Following curing of the helper plasmid, the strains were maintained in LB with 0.2% arabinose (Ara) at 30°C.

### Complementation titers and growth curve assays

For experiments in **Fig. 3**, CB4 and its derivatives were streaked out on LB-Ara-tet^10^ and grown overnight at 30°C. Single colonies were grown in 5 mL of LB-ara-tet10 overnight. The OD_600_ value of each culture was normalized to 1 before being serially 10-fold and plated on LB with either 0.2% arabinose (permissive), 0.2% glucose (non-permissive), 25 µM IPTG and 100 µM IPTG. Titer plates were incubated for 12-14 h at 37°C. For growth curves, OD-normalized cultures were diluted 1:1000 into ½ LB 0 NaCl with either 0.2% arabinose, 0.2% glucose or 25 µM IPTG and grown at 42°C, with data points measured every 5 min. Data presented are representative of three independent biological replicates. For growth curves shown in **SI Fig. 6**, TU122/pAV23 and the WT control (MG1655) were grown either under the same conditions as what was used for the sp-tracking experiments (see below), i.e., at 37°C in LB with or without 25 µM IPTG, or at 42°C in ½ LB 0 NaCl with or without 250 µM IPTG. The Halo-fusion of PBP1b rescued growth to WT levels in both conditions.

### Quantification of protein levels *in vivo* strains by western blotting

Individual colonies of complementation strains were grown overnight in LB-Ara-tet^10^ at 30°C. These cultures were diluted 1:1000 into 5 mL LB-Ara containing 25 µM of IPTG and cultured 30°C until OD600 = 0.5-0.8. Equivalent number of cells were centrifuged for each culture at 4,000 g for 10 min, pellets were washed once with ice-cold 1xPBS and flash frozen in liquid nitrogen. Each pellet was solubilized in 100 µL of solution composed of 1xSDS loading buffer and 50 mM HEPES pH 7.5, 150 mM NaCl, 2 mM MgCl_2_ and 0.25 units/µl benzonase and incubated at room temperature for at least 30 min. Following solubilization, the samples were centrifuged at 4,000 for 2 min to remove cell debris, and 10 µL were run on SDS-PAGE. The gel was transferred to a PVDF low fluorescence membrane using the Trans-Blot Turbo Transfer System (Bio-Rad). The membrane was blocked with 3% BSA 1xPBS for 1 h at room temperature, then incubated with primary antibody solution (rabbit α-LpoB antibody diluted 1:5,000 and mouse α-RpoA diluted 1:10,000, PMID: 21183074) for 1 h at room temperature. The membrane was then washed 3 times with 1xPBS, incubated with secondary antibody solution (IRDye 800CW Anti-Rabbit IgG Goat Secondary Antibody and IRDye 680RD Goat anti-Mouse IgG Secondary Antibody, LI-COR Biosciences) for 30 min at room temperature and washed again 3 times with 1xPBS. The membrane was visualized using a fluorescence imager (Amersham Typhoon 5).

Western blotting for sp-tracking strains was carried out as above, except that the strains were outgrown at 37°C in LB containing 0-1000 µM of IPTG. RpoA loading control was detected as above, and Halo-tagged PBP1b was detected using a mouse α-Halo tag monoclonal antibody (Promega).

### Sample preparation for single-particle-tracking experiments and data collection

Overnight cultures of TU122/pAV23 and TU122/pSI237 were diluted 1:1000 into 5 mL of LB supplemented with 25 µM IPTG and grown for 2-2.5 h until OD600 = 0.3-0.5. Janelia Fluor HaloTag ligand 554 (JFX554, CS315101, Promega) was added to cells at a working concentration of 20 nM and cells were grown for an additional 20 min. For each condition, 1 mL of cells was harvested by centrifugation (5,000 g for 2 min), washed three times with LB and resuspended in 25-50 µL of LB for TIRF fluorescence imaging.

Cells were added to high precision #1.5 coverslips (Corning, Millipore Sigma), placed on a 2% (w/v) agarose pad in LB and imaged at 37°C on a Nikon Ti inverted microscope equipped with a 561-laser line, a Plan Apo 100x/1.4 Oil Ph3 DM objective and a Andor Zyla 4.2 Plus sCMOS camera. Fluorescence time-lapse series were collected with 2×2 binning at 30 ms exposure time for 30 s and at 225 ms exposure time for 2 min, with a single brightfield reference image for each movie.

### Single-particle tracking analysis

Particle tracking was performed in Fiji using the TrackMate v7.11.1 plugin^71^. Spots were detected using the LoG-detector with an estimated object diameter of 0.4 µm and a quality threshold of 20. An intensity threshold on the brightfield signal was set to exclude any spurious fluorescent spots outside of the cell boundaries. Spots were linked using the LAP Tracker with a maximum linking distance of 0.4 µm and a maximum frame gap of 2. Tracks consisting of at least 4 spots were exported and further analyzed using the publicly available Spot-ON analysis platform^72^. To estimate the populations of immobile and freely diffusing molecules, tracks were fit to a two-state model. For all conditions the fit parameters were left free, to ensure unbiased estimates, with the exception of fixed cell controls where the lower bound on the fraction of the immobile molecules was fixed to 0.8 to allow the algorithm to converge. We used a custom Matlab script to analyze the exported track files and calculate the dwell time distributions and fits of the immobile states. Briefly, for each track containing at least 8 spots, trajectories of instantaneous jump distance as a function of time were calculated, and abrupt changes in the mean jump size were detected using a thresholding function. An immobile threshold of 70 nm for the maximum jump distance was set based on the single jump distance distribution of the fixed cell control (**SI Fig. 6d**). Periods of immobility were assigned to uncensored time segments lasting at least four frames, in which the average step size was maintained below the 70 nm threshold (**SI Fig. 6d**). Resulting histograms of time segments were fit to exponential distributions to extract values of dwell time. Single-particle tracking data collection parameters and statistics are summarized in **SI Table 4**.

---

**Figure S1:**
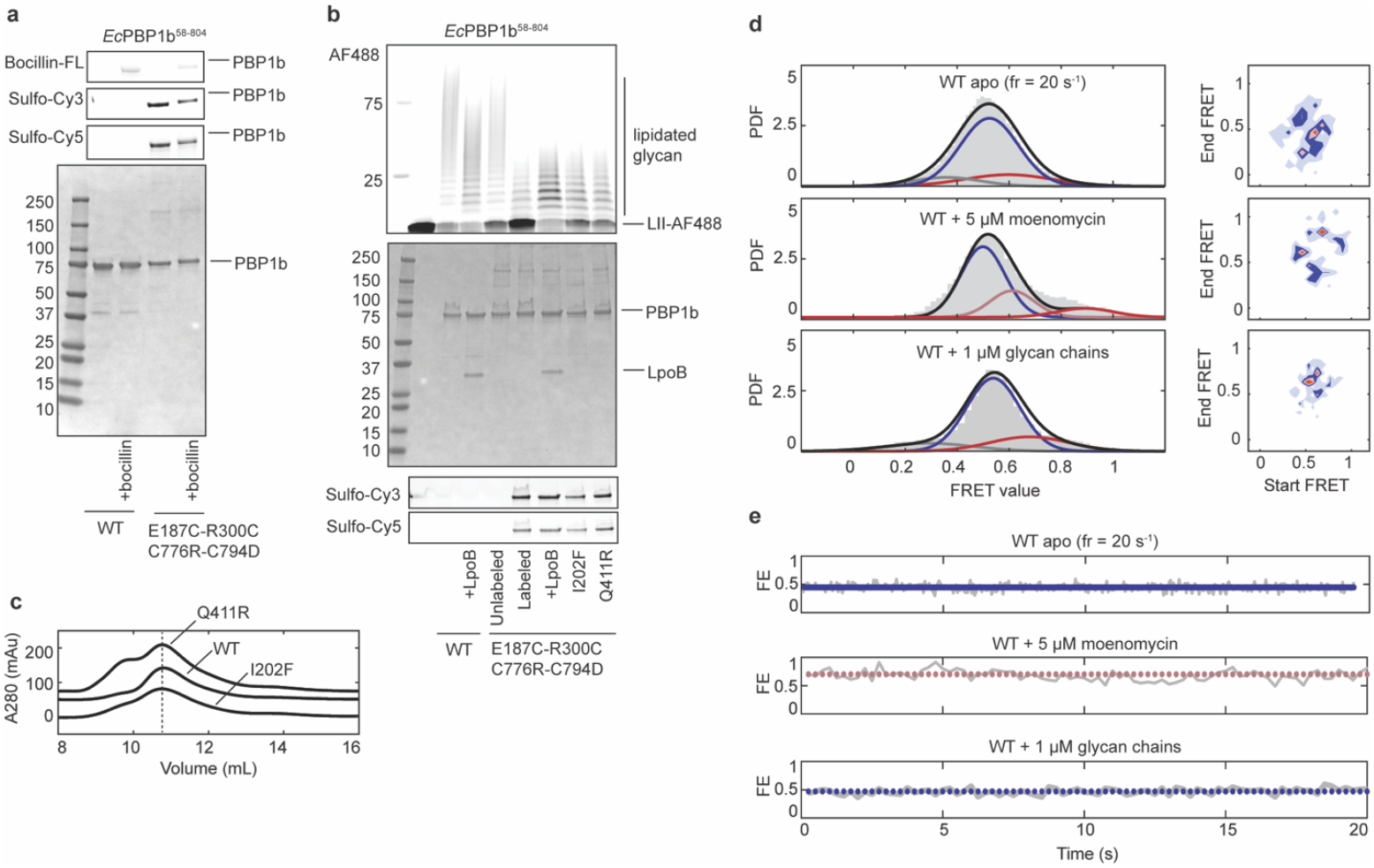
Biochemical validation of *Ec*PBP1b smFRET dynamics constructs. **a**, Coomassie, fluorescence and bocillin gels of the EcPBP1b smFRET imaging construct show that the TP active site retains bocillin binding in the absence of the structural disulfide and that engineered cysteines label specifically and efficiently with both Cy3 and Cy5 fluorophores. **b**, GT activity assay, coomassie and fluorescence gels for construct in (**a**) and suppressor variants (I202F, Q411R). PBP1b retains moderate GT activity after labeling and is robustly activated by LpoB and by suppressor mutations. **c**, Size-exclusion elution profiles for constructs in (**b**). **d**, PDF histograms of *Ec*PBP1b construct from (**a**) imaged at a fast frame rate (top), in the presence of moenomycin (middle), or with glycan chains (bottom), alongside transition density plots for each condition. **e**, Example single-molecule trajectories corresponding to population histograms in (**d**).

**Figure S2:**
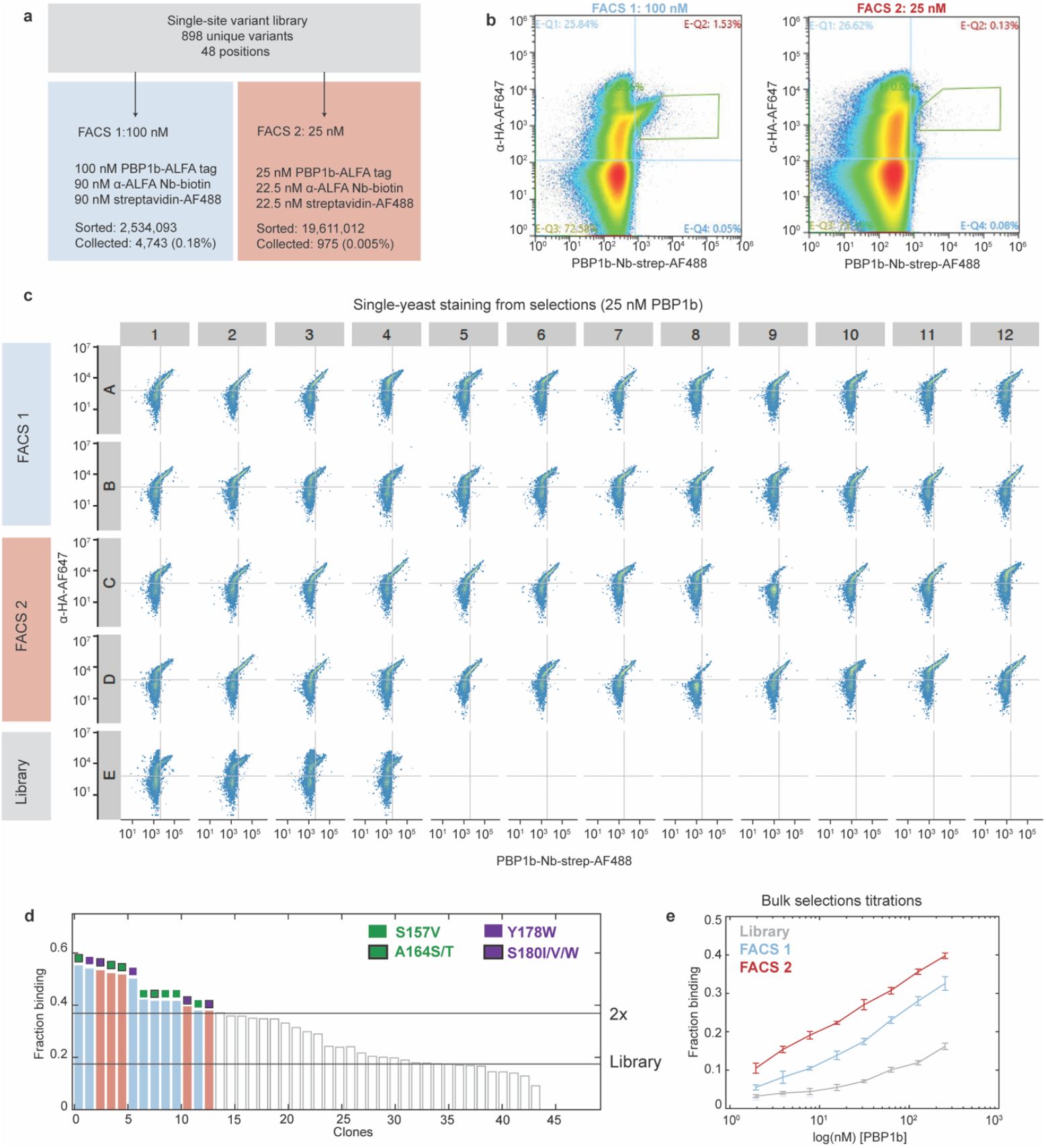
LpoB affinity maturation campaign. **a**, Flowchart of LpoB selections. Variants with improved affinity for PBP1b were selected through two rounds of fluorescence-activated cell sorting (FACS). **b**, FACS plots show gates on the yeast populations that were collected. **c**, Single-cell staining of individual yeast clones from the two rounds of FACS. **d**, Bar graph showing calculated bound fraction from yeast clones in (**c**). Clones that had at least 2-fold higher bound fraction than the naïve library were selected for subsequent sequencing and biochemical validation. Mutated positions are indicated for each clone. **e**, PBP1b titration curves with the two rounds of FACS, alongside the naïve yeast-display library control. FACS selections progressively improve bulk affinity of the isolated population for PBP1b.

**Figure S3:**
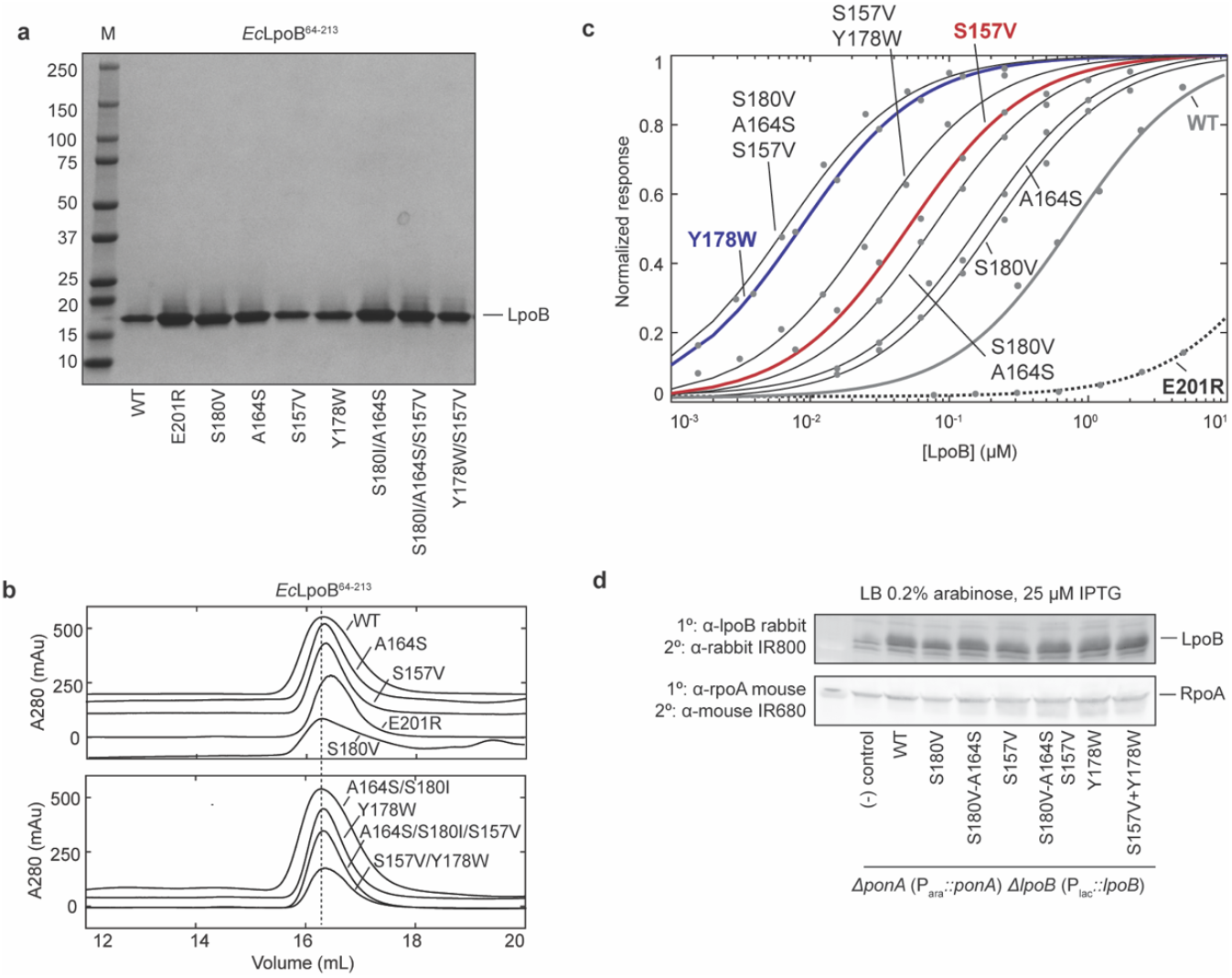
Affinity matured LpoB variants. **a**, Coomassie gel of engineered LpoB variants. **b**, Size-exclusion profiles of variants from (**a**) show a monodispersed elution profile for each construct. **c**, Example titration curves of LpoB variants from (**a**). Binding parameters and statistics are outlined in SI Table 2. **d**, Western blot analysis of the strains in Fig. 3 shows expression of all LpoB variants under conditions that match those of complementation assays.

**Figure S4:**
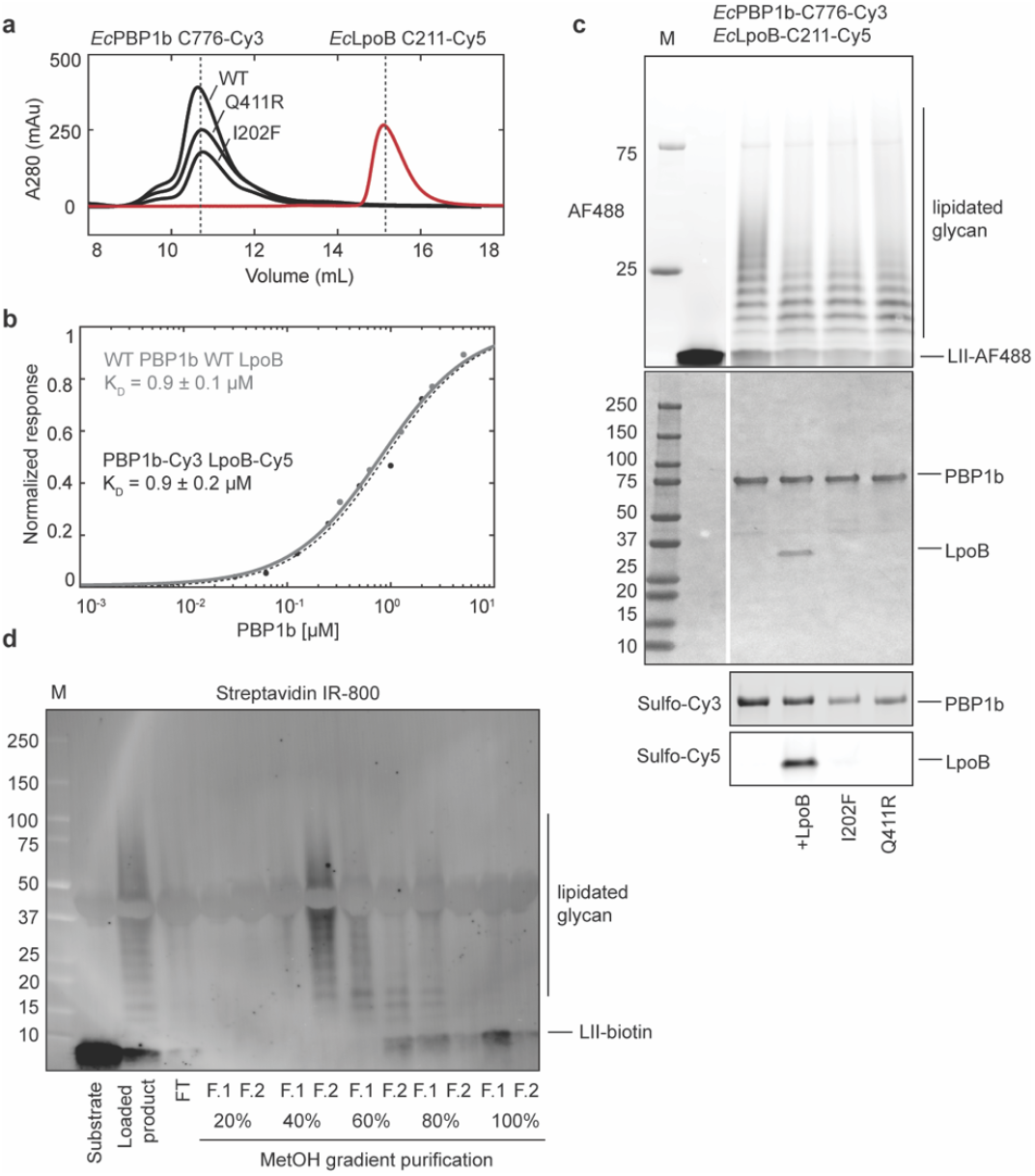
Biochemical validation of reagents used in smFRET binding assays. **a**, Size-exclusion runs *Ec*PBP1b and *Ec*LpoB variants for smFRET binding experiments show a monodispersed elution profile for each construct. **b**, BLI titration curve for the fluorescently labeled PBP1b and LpoB constructs from (**a**), compared to the WT PBP1b and LpoB control curve reproduced from SI Fig. 3c for convenience. Binding between PBP1b and LpoB is fully preserved post-labeling. **c**, GT assay, coomassie control and fluorescent gels of constructs from (**a**) show efficient labeling of all constructs and full retention of activity. Bypass mutants (I202F, Q411R) or LpoB addition exhibit the signature shift in the distribution of glycan chain lengths to shorter products, suggestive of increase in the rate of initiation. **d**, Anti-streptavidin blot shows the process of glycan chain purification. Polymerized product was loaded onto a C18 BakerBond column, washed and eluted with a methanol gradient to separate unpolymerized lipid II substrate from glycan chains. Glycan products in the resulting fractions were then biotinylated and detected by western blotting with Streptavidin IR-800 dye. Fractions 40% F.2 and 60% F.1 that were found to contain glycan chains without residual monomeric lipid II were used for experiments in Fig. 4.

**Figure S5:**
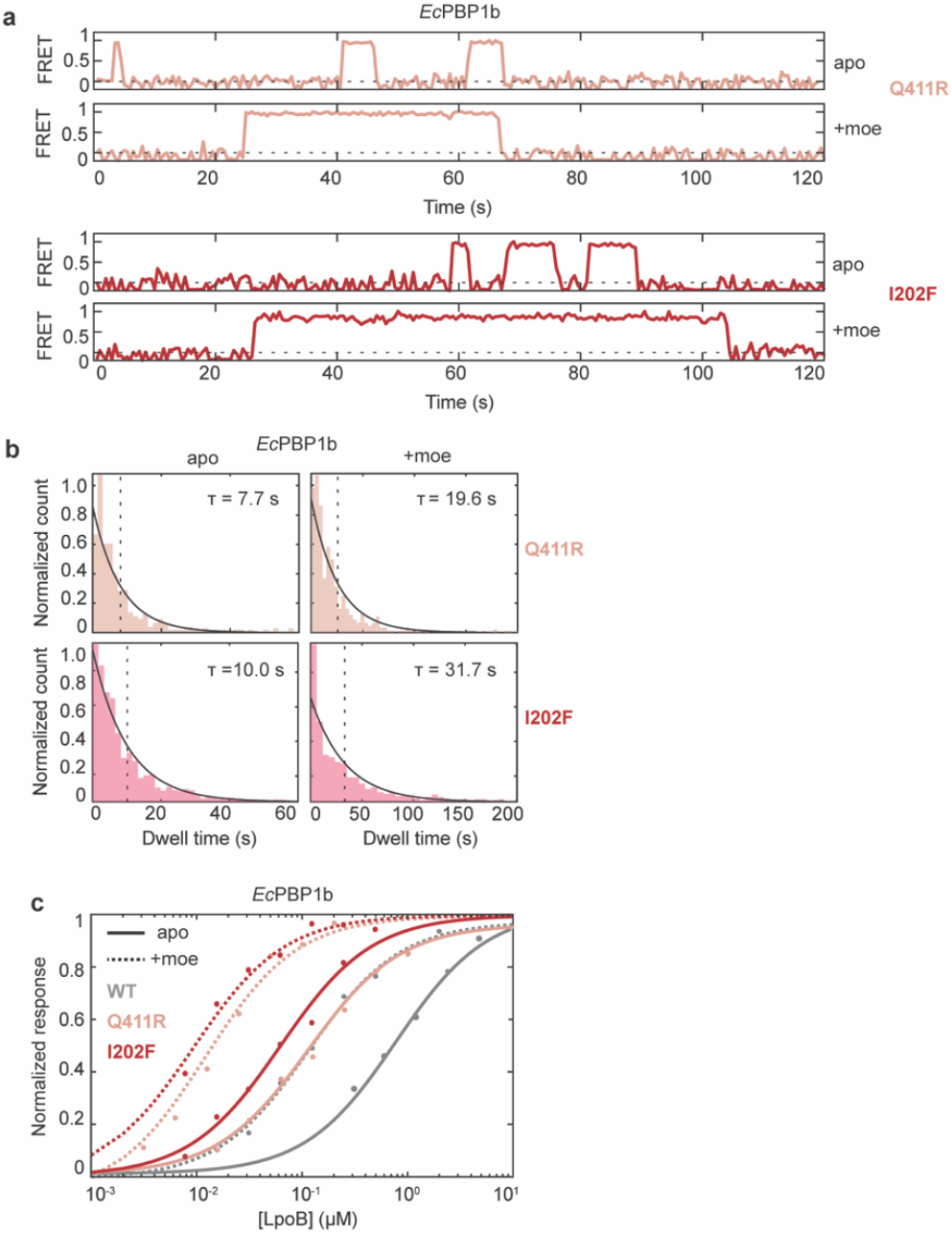
Suppressor variants increase LpoB affinity for PBP1b. **a**, Example trajectories of smFRET binding experiments with either Q411R or I202F PBP1b variants in the presence or absence of moenomycin. Suppressor mutations and substrate mimetic both increase the dwell time of the bound state. **b**, Dwell time histograms and fits of samples from (**a**). Full statistics are reported in SI Table 3. **c**, Example BLI binding curves showing that binding affinities determined through bulk and smFRET are similar. Full statistics are reported in SI Table 2.

**Figure S6:**
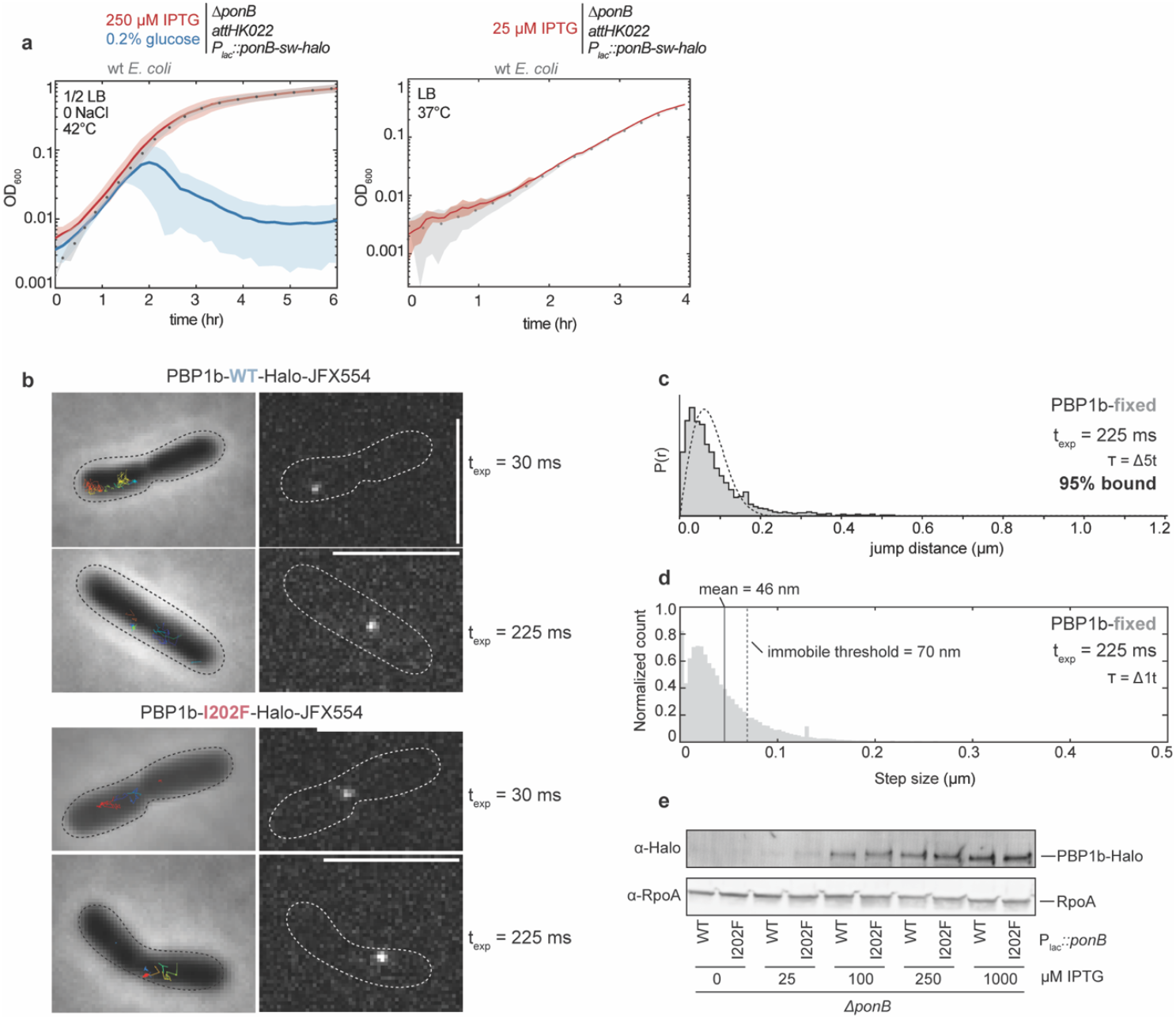
Single-particle experiments with Halo-tagged PBP1b WT and I202F. **a**, Growth curves of WT (MG1655) and Δ*ponB* P_*lac*_*::ponB-Halo-sw* fusion (TU122/pAV23) strains grown under non-permissive conditions (½ LB 0 NaCl at 42°C) or under standard conditions used for sp-tracking experiments (LB at 37°C). Upon induction, PBP1b-Halo strain (TU122/pAV23) shows similar growth rates as WT *E. coli* cells. **b**, Representative fluorescence micrographs (right) and corresponding brightfield images (left) with single-particle tracks overlaid of WT PBP1b-Halo (TU122/pAV23) and I202F PBP1b-Halo (TU122/pSI237) labeled with JFX554 fluorophore. **c**, Jump distance distribution and fit of fixed cells expressing PBP1b-Halo WT (TU122/pAV23), imaged with a long exposure time (225 ms). **d**, Step size distribution of dataset from (**c**) with the mean step size (46 nm) and the chosen immobile threshold (mean + ½ standard deviation = 70 nm) shown as solid and dotted lines, respectively. **e**, Western blot analysis shows expression levels of strains from (**b**) with 0-1000 μM IPTG induction.

**Figure S7:**
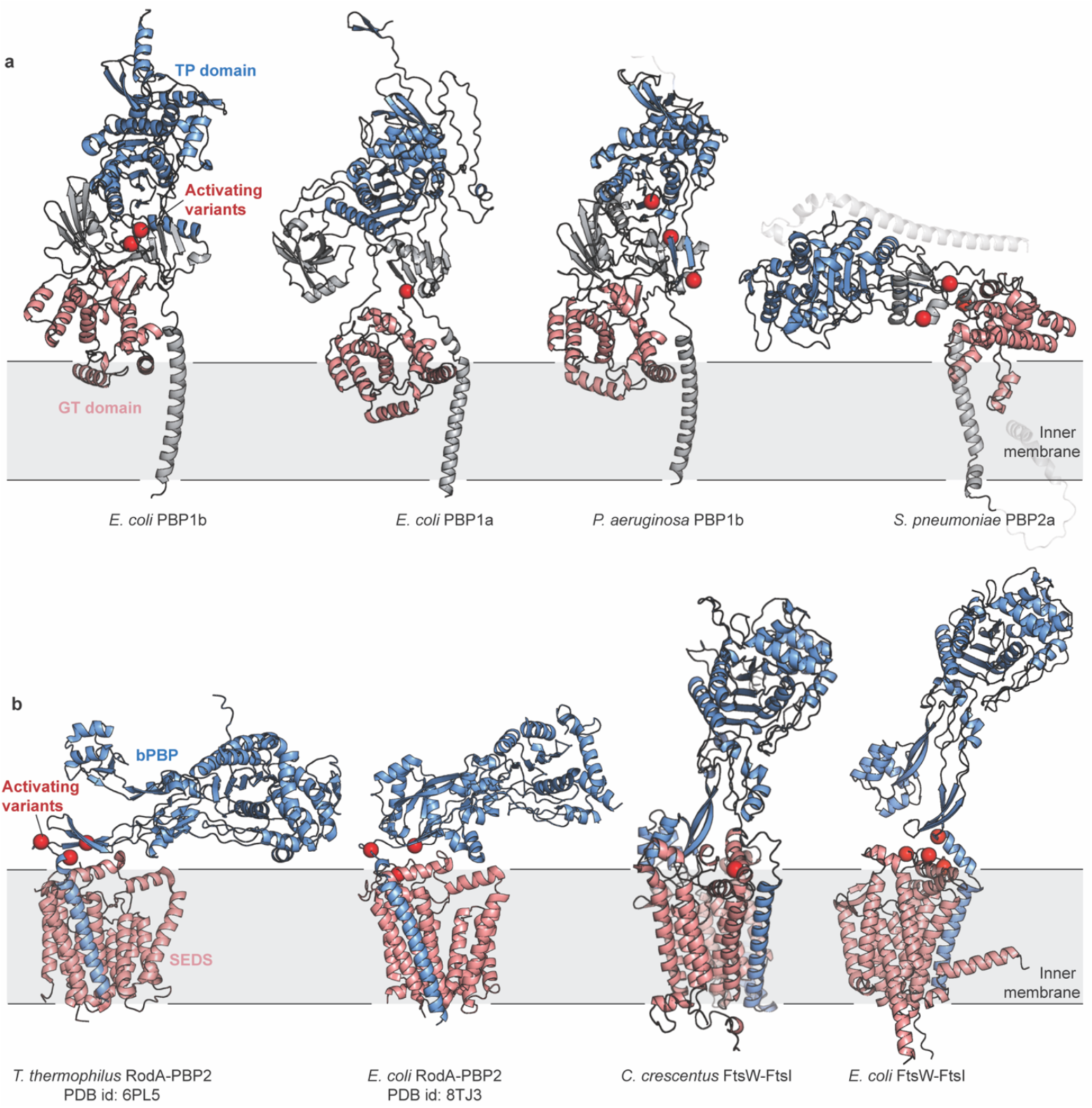
Activating mutations suggest conserved regulation in a diverse set of aPBP and SEDS-bPBP PG synthases. AlphaFold 3 models or structures of aPBP and SEDS-bPBP synthases in **a**, and **b**, respectively. GT (pink) and TP (blue) enzymatic domains are colored according to Uniprot domain assignment and activating mutations are shown as red spheres.

**Table S1:** smFRET dynamics data collection statistics.

| Condition | Exposure time (ms) | Total trajectories | Total data points | Mean FRET value (% occupancy) |
| --- | --- | --- | --- | --- |
| pSI228 | 250 | 344 | 59,555 | 0.31 (13%)<br>0.53 (70%)<br>0.65 (17%) |
| pSI228 | 50 | 191 | 59,703 | 0.35 (8%)<br>0.51 (80%)<br>0.62 (12%) |
| pSI228 + lipid II | 250 | 337 | 43,530 | 0.40 (8%)<br>0.61 (63%)<br>0.75 (29%) |
| pSI228 + moenomycin | 250 | 302 | 29,315 | 0.47 (56%)<br>0.62 (28%)<br>0.81 (16%) |
| pSI228 + glycan chains | 250 | 297 | 39,356 | 0.29 (12%)<br>0.53 (67%)<br>0.67 (19%) |
| pSI228 + LpoB | 250 | 505 | 93,111 | 0.44 (54%)<br>0.64 (27%)<br>0.83 (19%) |
| pSI228 + lipid II + LpoB | 250 | 205 | 32,684 | 0.46 (10%)<br>0.60 (62%)<br>0.70 (28%) |
| pSI229 | 250 | 354 | 75,451 | 0.55 (65%)<br>0.71 (35%) |
| pSI265 | 250 | 555 | 98,818 | 0.49 (51%)<br>0.65 (34%)<br>0.80 (15%) |
| pSI228 + pSI250 | 250 | 388 | 65,890 | 0.40 (62%)<br>0.65 (28%) |
| pSI228 + pSI296 | 250 | 244 | 35,100 | 0.36 (50%)<br>0.59 (31%)<br>0.79 (19%) |
| pSI228 + pSI299 | 250 | 286 | 41,396 | 0.37 (85%)<br>0.56 (15%) |
| pSI228 + pSI312 | 250 | 340 | 57,211 | 0.42 (71%)<br>0.66 (29%) |

**Table S2:** BLI binding parameters of LpoB and PBP1b variants.

| PBP1b variant | LpoB variant | $K_D$ (nM) | $k_{off}$ (s<sup>-1</sup>) |
| --- | --- | --- | --- |
| pSI199 | pSI259 | 900 ± 100 | 0.186 (0.189, 0.183) |
| pSI199 | pSI276 | >35 µM | NA |
| pSI199 | pSI298 | 300 ± 50 | 0.085 (0.086, 0.083) |
| pSI199 | pSI245 | 250 ± 50 | 0.074 (0.076, 0.073) |
| pSI199 | pSI296 | 60 ± 5 | 0.01 (0.011, 0.0099) |
| pSI199 | pSI294 | 12 ± 4 | 0.004 (0.0039, 0.0041) |
| pSI199 | pSI250 | 125 ± 25 | 0.035 (0.036, 0.034) |
| pSI199 | pSI299 | 10 ± 4 | 0.005 (0.0051, 0.0049) |
| pSI199 | pSI312 | 25 ± 3 | 0.0062 (0.0063, 0.0061) |
| pSI199 + moenomycin | pSI259 | 115 ± 10 | 0.048 (0.049, 0.047) |
| pSI200 | pSI259 | 111 ± 3 | 0.043 (0.044, 0.043) |
| pSI200 + moenomycin | pSI259 | 10 ± 4 | 0.003 (0.0031, 0.0029) |
| pSI201 | pSI259 | 53 ± 13 | 0.024 (0.025, 0.023) |
| pSI201 + moenomycin | pSI259 | 10 ± 5 | 0.004 (0.0043, 0.0041) |

**Table S3:** smFRET binding statistics.

| Condition | Exposure time (ms) | Total trajectories | Trajectories with binding events | $\tau_{\text{dwell}}$ (s) | $k_{\text{on}}$ ($\mu\text{M}^{-1}\text{s}^{-1}$) | Estimated $K_D$ (nM) |
| --- | --- | --- | --- | --- | --- | --- |
| pSI233 | 50 | 2511 | 1625 | 0.47 (0.44,0.50) | 2.3 (2.2, 2.6) | 915 |
| pSI233 + $\text{MnCl}_2$ | 50 | 1434 | 1005 | 0.53 (0.48,0.58) | 1.9 (1.85, 1.95) | 990 nM |
| pSI233 + lipid II | 100 | 795 | 630 | 9.0 (8.2,10.0) | 4.4 (4.3, 4.6) | 25 nM |
| pSI233 + lipid II + $\text{MnCl}_2$ | 250 | 1143 | 879 | 19.8 (18.5,21.2) | 3.2 (3.1, 3.3) | 15 nM |
| pSI233 + glycan | 50 | 2537 | 1701 | 0.70 (0.66,0.74) | 3.1 (3.0, 3.3) | 460 nM |
| pSI233 + glycan + $\text{MnCl}_2$ | 50 | 1716 | 977 | 2.45 (2.1,2.8) | 3.0 (3.1, 3.3) | 127 nM |
| pSI233 + moeno-mycin | 100 | 220 | 209 | 4.8 (4.3,5.4) | 4.6 (4.5, 4.8) | 45 nM |
| pSI234 | 250 | 660 | 635 | 10.0 (9.6,10.6) | 4.2 (4.1, 4.3) | 24 nM |
| pSI234 + moeno-mycin | 250 | 543 | 521 | 31.7 (29.6,34.0) | 2.5 (2.4, 2.6) | 12 nM |
| pSI242 | 250 | 837 | 596 | 7.7 (7.0,8.3) | 2.4 (2.3, 2.5) | 62 nM |
| pSI242 + moeno-mycin | 250 | 560 | 461 | 19.6 (18.0-21.4) | 2.5 (2.4, 2.6) | 20 nM |

**Table S4:** Single-particle tracking data collection.

| Condition | Exposure time (ms) | Total trajectories | $\tau_{\text{dwell}}$ (s) | Bound fraction |
| --- | --- | --- | --- | --- |
| PBP1b-Halo-sw | 30 | 2275 | NA | 0.47 |
| PBP1b-Halo-sw | 30 | 858 | NA | 0.53 |
| PBP1b-Halo-sw | 30 | 944 | NA | 0.57 |
| PBP1b-Halo-sw | 30 | 536 | NA | 0.62 |
| PBP1b-I220F-Halo-sw | 30 | 443 | NA | 0.65 |
| PBP1b-I220F-Halo-sw | 30 | 497 | NA | 0.78 |
| PBP1b-I220F-Halo-sw | 30 | 640 | NA | 0.75 |
| PBP1b-I220F-Halo-sw | 30 | 656 | NA | 0.73 |
| PBP1b-Halo-sw | 225 | 5197 | 4.8 | NA |
| PBP1b-I220F-Halo-sw | 225 | 7361 | 4.8 | NA |

**Table S5:** In vitro plasmids.

| Plasmid | Genotype | Source |
| --- | --- | --- |
| pSI191 | ColA-T7:: His6-SUMO-Flag-ALFA- *EcPBP1b*(58-804) | This study |
| pSI228 | ColA-T7:: His6-SUMO-Flag-ALFA- *EcPBP1b*(58-804) E187C-R300C-C776R-C794D | This study |
| pSI229 | ColA-T7:: His6-SUMO-Flag-ALFA- *EcPBP1b*(58-804) E187C-R300C-C776R-C794D-I202F | This study |
| pSI265 | ColA-T7:: His6-SUMO-Flag-ALFA- *EcPBP1b*(58-804) E187C-R300C-C776R-C794D-Q411R | This study |
| pCB39 | *bla* T7::His6-*EcLpoB*(21-213) | Paradis-Bleau *et al.* 2010 |
| pSI204 | *bla* T7::His6-*EcLpoB*(21-213)-S211C | This study |
| pSI233 | ColA-T7:: His6-SUMO-Flag-ALFA- *EcPBP1b*(58-804) C794A | This study |
| pSI234 | ColA-T7:: His6-SUMO-Flag-ALFA- *EcPBP1b*(58-804) I202F-C794A |  |
| pSI242 | ColA-T7:: His6-SUMO-Flag-ALFA- *EcPBP1b*(58-804) Q411R-C794A |  |
| pSI199 | ColA-T7:: *lpp*-C*-*EcPBP1b*(82-804)-ALFA-PrtC | This study |
| pSI200 | ColA-T7:: *lpp*-C*-*EcPBP1b*(82-804)-Q411R-ALFA-PrtC | This study |
| pSI201 | ColA-T7:: *lpp*-C*-*EcPBP1b*(82-804)-I202F-ALFA-PrtC | This study |
| pSI259 | ColA-T7::PrtC-*EcLpoB*(64-213) | This study |
| pSI276 | ColA-T7::PrtC-*EcLpoB*(64-213)-S153A-S157A- E201R | This study |
| pSI298 | ColA-T7::PrtC-*EcLpoB*(64-213)-S180V | This study |
| pSI245 | ColA-T7::PrtC-*EcLpoB*(64-213)-A164S | This study |
| pSI296 | ColA-T7::PrtC-*EcLpoB*(64-213)-S157V | This study |
| pSI294 | ColA-T7::PrtC-*EcLpoB*(64-213)-Y178W | This study |
| pSI250 | ColA-T7::PrtC-*EcLpoB*(64-213)-A164S-S180I | This study |
| pSI299 | ColA-T7::PrtC-*EcLpoB*(64-213)- A164S-S180I-S157V | This study |
| pSI312 | ColA-T7::PrtC-*EcLpoB*(64-213)-Y178-S157V | This study |

**Table S6:** In vivo plasmids.

| Plasmid | Genotype | Source |
| --- | --- | --- |
| pAV23 | attHK022 P<sub>lac</sub>::halo-ponB-sw (PBP1b-gamma) | This study |
| pSI237 | attHK022 P<sub>lac</sub>::halo-ponB-I202F-sw (PBP1b-gamma) | This study |
| pTB309 | attI P<sub>ara</sub>::ponA | Paradis-Bleau *et al.* 2010 |
| pSI241 | attHK022 P<sub>lac</sub>::lpoB | This study |
| pSI280 | attHK022 P<sub>lac</sub>::lpoB-S153A-S157A-E201R | This study |
| pSI258 | attHK022 P<sub>lac</sub>::lpoB-A164S-S180I | This study |
| pSI300 | attHK022 P<sub>lac</sub>::lpoB-Y178W | This study |
| pSI302 | attHK022 P<sub>lac</sub>::lpoB-S157V | This study |
| pSI304 | attHK022 P<sub>lac</sub>::lpoB-S180V | This study |
| pSI305 | attHK022 P<sub>lac</sub>::lpoB-A164S-S180I-S157V | This study |
| pSI314 | attHK022 P<sub>lac</sub>::lpoB-Y178W-S157V | This study |

**Table S7:** In vivo strains.

| Strain | Genotype | Source |
| --- | --- | --- |
| TU122/ pAV23 | MG1655 ΔlacIZYA<>frt mrcB(ponB)<>frt attk022 pHC949 | This study |
| TU122/pSI237 | MG1655 ΔlacIZYA<>frt mrcB(ponB)<>frt attk022 pSI260 | This study |
| CB4 | MG1655 ΔlacIZYA<>frt mrcA(ponA)<>frt attlTB309 ycfM::kan | Paradis-Bleau *et al.* 2010 |
| CB4/pSI241 | MG1655 ΔlacIZYA<>frt mrcA(ponA)<>frt attlTB309 attk022 pSI241 ycfM::kan | This study |
| CB4/pSI280 | MG1655 ΔlacIZYA<>frt mrcA(ponA)<>frt attlTB309 attk022 pSI280 ycfM::kan | This study |
| CB4/pSI258 | MG1655 ΔlacIZYA<>frt mrcA(ponA)<>frt attlTB309 attk022 pSI258 ycfM::kan | This study |
| CB4/pSI300 | MG1655 ΔlacIZYA<>frt mrcA(ponA)<>frt attlTB309 attk022 pSI300 ycfM::kan | This study |
| CB4/pSI302 | MG1655 ΔlacIZYA<>frt mrcA(ponA)<>frt attlTB309 attk022 pSI302 ycfM::kan | This study |
| CB4/pSI304 | MG1655 ΔlacIZYA<>frt mrcA(ponA)<>frt attlTB309 attk022 pSI304 ycfM::kan | This study |
| CB4/pSI305 | MG1655 ΔlacIZYA<>frt mrcA(ponA)<>frt attlTB309 attk022 pSI305 ycfM::kan | This study |
| CB4/pSI314 | MG1655 ΔlacIZYA<>frt mrcA(ponA)<>frt attlTB309 attk022 pSI314 ycfM::kan | This study |

## Notes

### Competing Interest Statement

A.C.K. is a cofounder and consultant for biotechnology companies Tec-tonic Therapeutic and Seismic Therapeutic, and for the Institute for Protein Innovation, a non-profit research institute.

